# Gene expression analysis of carbohydrate catabolism in *Leucoagaricus gongylophorus* LEU18496

**DOI:** 10.1101/2025.03.21.644473

**Authors:** Freddy Castillo-Alfonso, Gabriela Cejas-Añón, Cecilio Valadez-Cano, Juan Carlos Sigala Alanis, Gabriel Vigueras-Ramírez, Roberto Olivares-Hernández

## Abstract

Transcriptomic profiles were obtained by RNA-seq of *Leucoagaricus gongylophorus* growing on two carbon sources (glucose and cellulose) during different growth phases (exponential and stationary) with the aim of analyzing the variations in metabolism depending on the substrate and/or growth state. Principal component analysis (PCA) showed that 99.5% of the variation in gene expression is related to the substrate (PC1), clearly separating the experimental conditions, while the differences between growth phases are reflected in secondary components (PC2). Special attention was paid to the differential expression of CAZymes and FOLymes enzymes, demonstrating that, during the exponential phase of cellulose culture, there is an overexpression of enzymes from the GH6 and GH7 families (cellulases and cellobiosehydrolases), related to the degradation of complex polysaccharides such as cellulose. During the exponential phase of growth on glucose, expression of CAZymes enzymes (*β*-glucosidase, pectinase and endo-*β*-1,3-glucanase) was observed, in addition to an overexpression of laccases in this carbon source. Enrichment analysis identified significantly enriched functions related to carbohydrate catabolism and transport, evidence of the metabolic versatility of *L. gongylophorus*. High expression of the *creA* repressor factor was found in the presence of glucose, suggesting its regulatory role in the modulation of carbohydrate degradation. In the stationary phase, genes related to the response to oxidative stress and nutrient solubilization were activated under glucose, in contrast to cellulose, where the activation of secondary metabolism pathways was promoted, including the overexpression of trichodiene synthase.

## 1. Introduction

*Leucoagaricus gongylophorus* Möller Singer (Agaricales, Agaricaceae) is a basidiomycete able to establish mutualistic relationships with ants of the genera *Atta* and *Acromyrmex*. Ants build a nest made of lignocellulosic plant material which is used as a substrate by fungi to build their gardens. This organism is capable of producing a wide range of enzymes that degrade complex polysaccharides of plant material, converting this organic matter into a substrate that can be assimilated by ants [3]

The genome of *L. gongylophorus* LEU18496 (121 Mbp), mutualistic fungus with *Atta mexicana* ants, was reported to have 6,748 genes and a genome structure with bimodal distribution of GC content. A high relative abundance of CAZymes (carbohydrate-active enzymes) and FOLymes (lignin oxidative enzymes) was also reported for this genome [9]. The fungus has been successfully isolated and identified from the fungal gardens of *A. mexicana* and several phylogenetic analyses confirming its identity. The production of CAZymes by *L. gongylophorus* is central to this symbiosis, enabling the ants to metabolize cellulose and produce sugars consumed by the colony [31]. *L. gongylophorus* has been shown to thrive in laboratory cultures using glucose as the unique carbon source demonstrating its versatility in utilizing different carbon substrates [49]. Previously, the identification of crucial genes related to the metabolism of complex carbohydrates such as plant polymers degraded by various CAZymes families had been reported [42]. These enzymes include glycosyl hydrolases (GHs), which act synergistically to break down cellulose, and oxidative enzymes like fungal oxidative lignin enzymes (FOLymes), which play a role in lignocellulose degradation. Understanding the regulation of CAZymes is critical, especially in industrial applications where efficient enzyme production is needed for biomass conversion processes [22].

In 2022, Conlon et al. used ^13^C isotopic labeling to analyze cellulose consumption in *L. gongylophorus* cultivars and the ant *A. colombica*, glucose consumption was used as a reference condition in the comparisons [11]. These researchers demonstrated the production of cellulases in potato dextrose agar medium without cellulose, evidencing the production of CAZymes in medium with glucose. They also compared gene expression using transcriptomes reported by Nygaard *et al*. in 2016 for *A. colombica* cultivars [33]. Leal-Dutra *et al*. in 2023 obtained transcriptomes of *L. gongylophorus* growing in potato dextrose medium for 30 days. They subsequently performed differential gene expression analysis, demonstrating the mechanisms of positive regulation in genes involved in autophagy and cell differentiation [25].

Gene expression can be regulated in fungi by various mechanisms such as carbon catabolic repression (CCR). This mechanism is a system that, in the presence of a monosaccharide such as glucose, inhibits the consumption of complex substrates such as cellulose. In filamentous fungi, the presence of glucose usually inhibits the synthesis of enzymes related to the degradation of lignocellulosic compounds. In this key mechanism, regulatory elements such as transcription factors, transposable elements, and chaperone proteins are involved [13]. Knowledge of these regulatory mechanisms may be essential to improve/optimize the industrial production of CAZymes and FOLymes, since circumventing the CCR may lead to higher enzyme productivity scenarios when these organisms are grown under less favored carbon sources.

The main goal of this study is to analyze the gene expression profiles of *L. gongylophorus* LEU18496 during exponential and stationary growth phases with glucose or cellulose as carbon sources. By examining the genes involved in polysaccharide metabolism, this research aims to elucidate how growth phases and substrate presence can influence enzyme production and the regulation of CAZymes and FOLymes. The analyses carried out in this research may contribute to deepening the knowledge of the regulatory elements in the production of CAZymes and FOLymes and attempt to diversify the biotechnological applications of the production of these enzymes in bioconversion processes of lignocellulosic materials or degradation of compounds of industrial interest.

## 2. Materials and methods

### 2.1. Cell culture and propagation

*L. gongylophorus* LEU18496 cultures were cultivated with different carbon sources to obtain biomass. The strain was propagated on malt extract agar (MEA-LP) containing 20 g/L of malt extract (CAS 8002-48-0), 5 g/L of bacteriological peptone (CAS 91079-38-8), 2 g/L of yeast extract (CAS 8013-01-2) and agar–agar (CAS 9002-18-0), and the pH was adjusted to 5.0. The cultures were incubated at 27 ± 1 °C. From these solid plate cultures of *L. gongylophorus*, 500 mL Erlenmeyer flasks with 120 mL of malt extract medium (CAS 8002-48-0) were inoculated and incubated for 15 days at 27 ± 1 °C and 120 rpm in an orbital shaker (New Brunswick I26 Incubator Shaker). The precultures were centrifuged aseptically and inoculated in 250 mL Erlenmeyer flasks with 80 mL of minimum salt medium [15] supplemented with glucose (CAS 50-99-7) or carboxymethylcellulose (CMC; CAS No. 9000-11-7) at a concentration of 10 g/L and incubated for 35 days at 27 ± 1 °C and 120 rpm in an orbital shaker (New Brunswick Scientific I26 Incubator Shaker). CMC is a cellulose derivative chemically modified by the introduction of carboxymethyl groups, which gives it water solubility.

To obtain biomass, samples were collected on days 21 and 35 of culture for each carbon source, resulting in a total of 4 samples. Cell cultures were centrifuged and frozen in a mixture of dry ice and ethanol for subsequent RNA extraction. A second experiment was conducted under the same conditions and using the same culture media, with the aim of obtaining biological replicates for each condition. This resulted in a total of 8 samples, including two biological replicates of growth in glucose and two biological replicates of growth in cellulose, sampled during the exponential and stationary phases of growth.

### 2.2. RNA extraction and sequencing

Extraction, library preparation, and RNA sequencing were performed by the Sequencing and Bioinformatics Information System (UNAM) sequencing service via the Illumina NextSeq platform. Briefly, RNA extraction was performed via TRIzol Reagent using 50 mg of biomass following manufacturer’s instructions. The sequencing process was performed using an Illumina NextSeq 500 platform, in a configuration of 2×76 cycles (paired end 150). A total of 1.5 mL of the solution with the libraries at a concentration of 1.8 pM was loaded into the sequencing equipment.

The Table 1 presents the details of the samples collected during the experiment. It includes the sample ID, the carbon source used (glucose or cellulose), the day of collection (either day 21 or day 35), the growth phase (exponential or stationary), the biological replicate number, and the RNA extraction process for each sample.

**Table 1.**
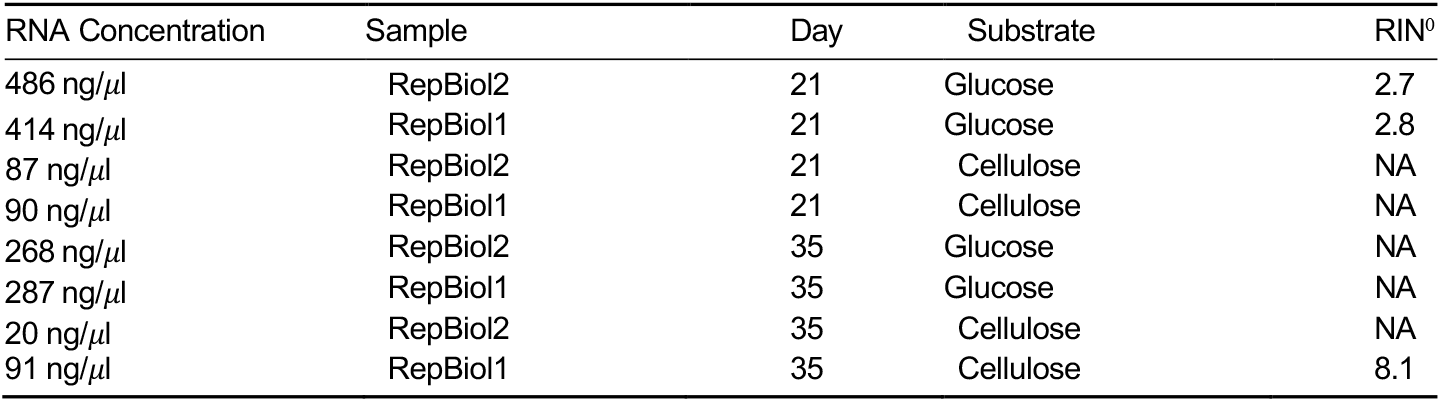
Technical parameters including RNA concentration, sample ID/biological replicate, collection day, substrate, and RNA Integrity Number (RIN).

| RNA Concentration | Sample | Day | Substrate | RIN<sup>0</sup> |
| --- | --- | --- | --- | --- |
| 486 ng/μL | RepBiol2 | 21 | Glucose | 2.7 |
| 414 ng/μL | RepBiol1 | 21 | Glucose | 2.8 |
| 87 ng/μL | RepBiol2 | 21 | Cellulose | NA |
| 90 ng/μL | RepBiol1 | 21 | Cellulose | NA |
| 268 ng/μL | RepBiol2 | 35 | Glucose | NA |
| 287 ng/μL | RepBiol1 | 35 | Glucose | NA |
| 20 ng/μL | RepBiol2 | 35 | Cellulose | NA |
| 91 ng/μL | RepBiol1 | 35 | Cellulose | 8.1 |

A total of eight libraries were obtained with an average of 10 million reads for each one. The quality of the data was analyzed via FastQC v 0.11.5 [6] to subsequently subject the raw data to a cleaning process (trimming) to remove sequencing adapters and low-quality bases via FASTp v 0.23.4 [10]. After cleaning, the data were analyzed again using FastQC v 0.11.5 to evaluate the results of the trimming process. Secondary metabolite gene clusters (BCG) were predicted using antiSMASH version 7.0.1 [4] with default parameters and via the *L. gongylophorus* reference genome (GCA022457215.3).

### 2.3. Transcriptome mapping and count table creation

The data obtained from RNA-seq were mapped via the *L. gongylophorus* reference genome (GCA022457215.3) [9]. The STAR program was used to align the sequencing reads with the reference genome once the genome indices were created. The genome index in STAR is a set of precomputed data structures representing the reference genome [12]. The HTseq tool [53] was used to generate tables of gene counts, given one or several SAM/BAM files with alignments and an annotation file with gene prediction features of the genomic assembly used as a reference [9]. Differential expression analysis was performed using DESeq2 [29], a statistical package in R used to analyze gene expression data.

The analysis was performed using glucose as the reference carbon source and the exponential phase as the reference growth phase. Quantile normalization was applied to ensure that the expression distributions of the genes were similar between different samples, adjusting for any systematic biases in the data. For differential expression analysis, the experimental design included two carbon sources, Cellulose and Glucose, and two growth phases, Exponential (Exp) and Stationary (Sta). The experimental conditions were defined as a factor variable (condition), which was structured as a combination of the carbon source and growth phase, with two biological replicates for each condition. The phenotype factor indicated the growth phase, with each phase repeated four times. The final design consisted of a total of 8 samples, representing the different combinations of carbon source, growth phase, and biological replicates. A principal component analysis (PCA) and a clustering sample heat map was performed using DESeq2 [29].

The statistical parameters for determining differential expression include a padj ≤ 1 × 10^−16^, indicating highly significant differences between conditions. Additionally, gene expression changes were evaluated using log_2_FoldChange (log_2_FC), calculated as the base-2 logarithm of the ratio between gene expression in the treatment and control conditions (glucose and exponential phase):

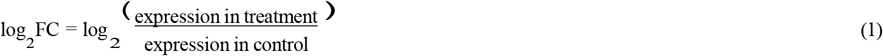

A log_2_FC > 1 was interpreted as a significant increase in gene expression, while log_2_FC < -1 indicated a significant decrease in expression in the experimental condition compared to the control (glucose and exponential phase).

## 3. Results and Discussion

### 3.1. Count tables and clustering

Transcriptional variations of *L. gongylophorus* LEU18496 growing on glucose or cellulose in exponential and stationary growth phases were investigated. Between 91% and 93% of the reads for all analyzed samples were mapped (Table S1). These alignment values demonstrate the quality of the samples processed to obtain the counting tables [24][37].

To determine the similarity and consistency of the transcriptomes obtained, we carried out a principal component analysis (PCA) of the normalized counts for each carbon source/phase of growth/biological replicate (Figure 1). The PCA graphs show a large grouping of the biological replicates, demonstrating the high degree of similarity in the intra-biological duplicate counts. In PC1, which maximizes the direction in which the observations vary the most [16], the highest percent of variance is found, thus allowing us to determine that there is a differential relationship between the variables analyzed. Except for the samples corresponding to the biological duplicate of *L. gongylophorus* growing on cellulose during the exponential phase, which presents the greatest variability in PC2 with a value around 12%, even so the greatest variance in PC1 dominates the separation between the compared samples. Based on the results presented in Figure 1, there is a differential count of expression between glucose and cellulose conditions in the two growth phases. Fundamentally when glucose and cellulose are compared in the exponential phase where there is a 99.5% variance in PC1, indicating that genetic regulation is broadly related to the substrate that is used as a carbon source. A lesser impact can be seen between the different phases of growth, although significant differences are also seen. The distribution of the samples was also verified using sample matrix distance, corroborating the before mentioned distribution where the samples corresponding to cellulose have the greatest location and the least grouping when they are clustered in a dendrogram that includes all the samples, see Figure S1. The results shown in Figure S1 confirm the differences observed in the PCA analysis, highlighting that the carbon source confers more variability in gene expression under the conditions analyzed. These results reinforce that gene expression is highly related to the carbon source, reflecting possible differential expression patterns according to the available substrate.

**Figure 1:**
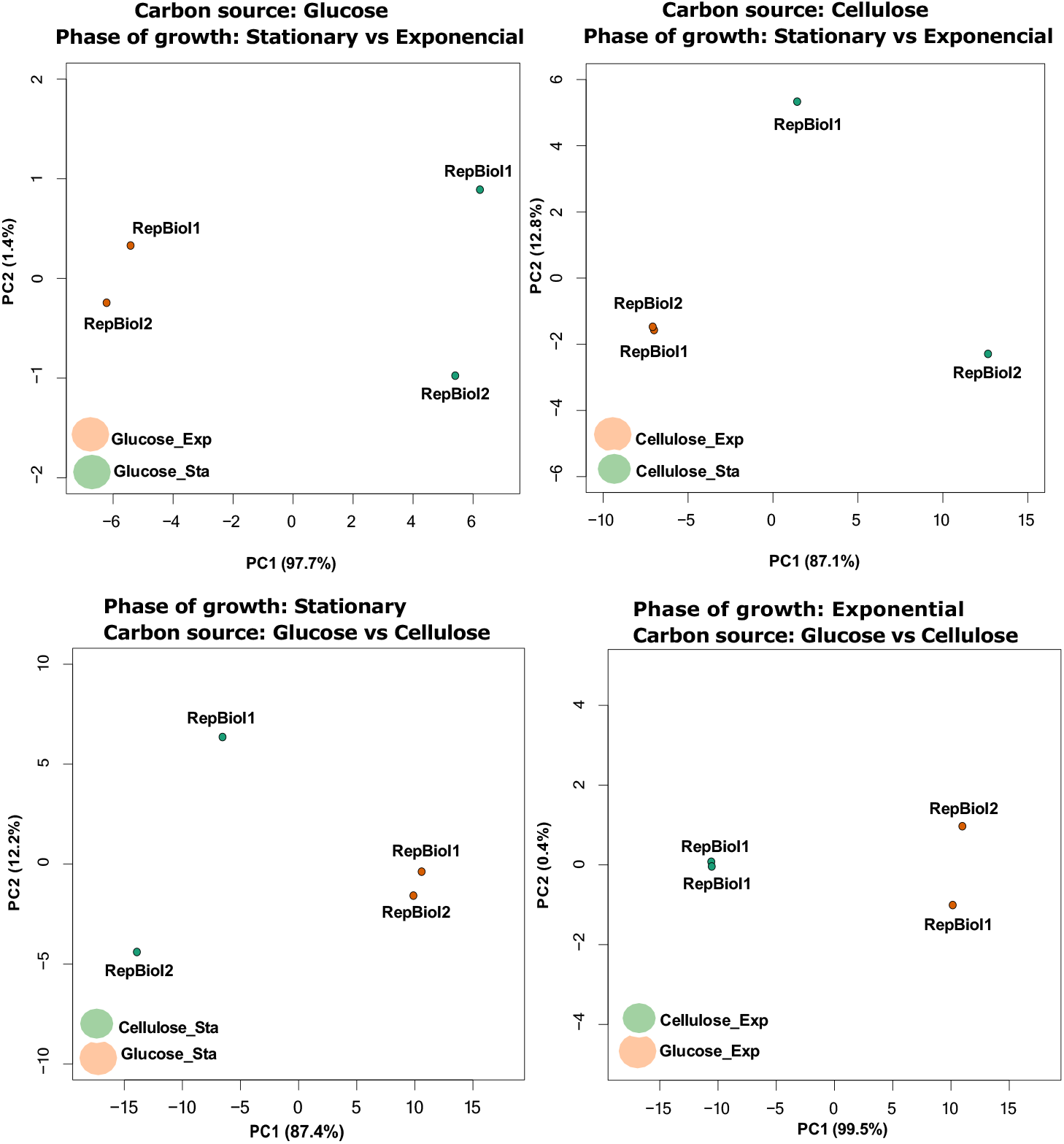
Principal component analysis (PCA) of the normalized counts obtained for each sample analyzed. The terms RepBiol correspond to the biological duplicates, glucose and cellulose are the conditions explored and Exp (Exponential phase) and Sta (Stationary phase) with the sampling times carried out and referred to as phenotypes in the analysis.

### 3.2. General overall expression

The transcriptomic gene expression levels of *L. gongylophorus* LEU18496 were analyzed across five functional categories: CAZymes, FOLymes, transporters and transcription factors (TFs). These expression levels were calculated based on a quartile-based analysis of transcriptomic data, providing a robust assessment of gene activity under the studied conditions (Table 2). The genomic assemblies reported show a high presence of CAZymes and FOLymes in the metabolism of *L. gongylophorus* [9]. Of the total CAZymes and FOLymes reported in the genome, 72.91% and 84% were detected in the transcriptomes, respectively, which shows the importance of the expression of these enzymes for *L. gongylophorus* as part of its life cycle (Table 2). Zhao et al., in 2013, used comparative analysis between different fungi belonging to different trophic groups and reported a large number of CAZymes in these genomes, species-specific, and plant-pathogenic fungi [57]. The abundance of this enzymatic group has also been reported for endophytic fungi [23]. Differential regulation in CAZymes related to lignocellulose catabolism has also been reported for basidiomycete fungi [40]

**Table 2.**
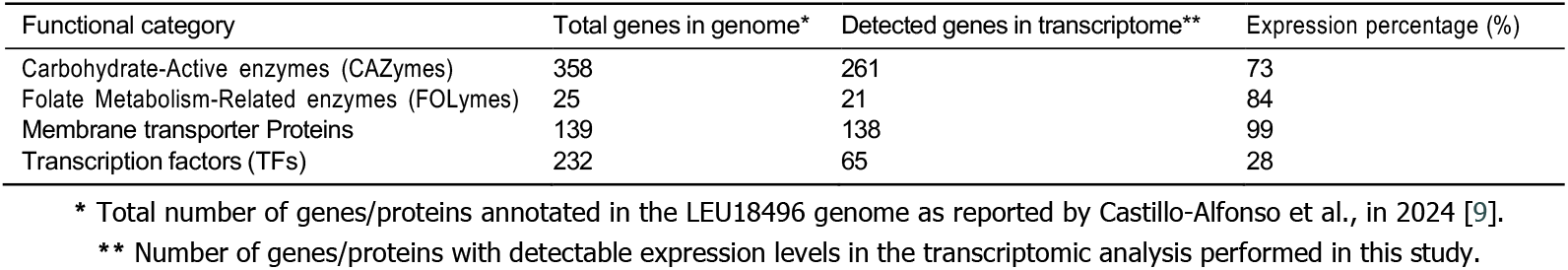
Functional categorization of genes in the *L. gongylophorus* LEU18496 reported genome and their expression in the transcriptome.

As the products of complex polysaccharide degradation must be internalized to satisfy cellular metabolism, we found that 99.3% of the different transporter families reported in the genome showed the highest correlation rate (Table 2). This phenomenon may be directly associated with the high availability of dissolved sugars related to cellulose degradation. The sugar transporter family (TC 2.A.1.1) are uniporter sugar transporters present in the fungi. These transporters participate in the assimilation of hexoses; as a result of cellulose degradation, various sugars are released into the environment, so the higher expression of these transporters in cellulose as a carbon source is marked by the higher availability of sugars for assimilation. When grown on Avicel (a mixture of microcrystalline cellulose and sodium carboxymethylcellulose), Neurospora crassa, a model organism for cellulosic material degradation, overexpresses transporters belonging to the MFS superfamily (major facilitator superfamily) [47]. In 2018, Nogueira et al. demonstrated the influence of hexose transporters on metabolic signaling and the expression of genes with cellulase functions [32]. Transcription factors (TF) presented the lowest number of genes founded versus the high number reported for the *L. gongylophorus* genome, with 28.0%. By detecting this level of expression, we can conclude that the conditions evaluated do not promote the activation of a large number of transcription factors. However, several transcription factors were found during the analyses that we will describe below. The CCAAT-binding complex (CBC) is a highly conserved heterotrimeric transcription factor (Hap) in fungi and regulates the transition between fungal hyphal growth and asexual reproduction. Furthermore, the Hap2/3/5 complex is involved in regulating cellulase expression in *Trichoderma reesei*, together with other regulators, such as *xyr1* and *ace2* [46]. The velvet transcription factor complex controls the rate of asexual or sexual development in response to light, promoting sexual development in the dark while stimulating asexual sporulation under illumination [28]. Transcription factors associated with sexual reproduction have been identified, and the taxonomic classification of *Leucoagaricus sp*. has been complex because ants have historically been shown to suppress sexual reproduction when growing in symbiosis [41]. On the other hand, the presence of sexual structures has only been reported under laboratory conditions and during the development of axenic cultures without the presence of ants. We found expression of the gene g5255 (Zinc finger C2H2 type), ortholog of the *creA* gene in all the conditions previously evaluated, a factor involved in carbon catabolic repression (CCR) in filamentous fungi. *creA* gene presents a higher expression in glucose, which coincides with its repressive function on genes that encode CAZymes. Later in the document we will delve into the expression of *creA*. The results derived from the expression analysis by quartiles reflect that there is a high presence of CAZymes and FOLymes that, when degrading the substrate, activate a large number of sugar transporters, while the transcription factors expressed suggest mechanisms for expressing enzymes in the presence of glucose and also give indications of sexual reproduction in *L. gongylophorus* during exponential phase of growth.

The high number of CAZymes and FOLymes expressed in the transcriptomes of *L. gongylophorus*, due to the central role of complex polysaccharide degradation as part of its life cycle and the expression of transporters to internalize degraded sugars, demonstrates the high specialization of this fungus to satisfy cellular metabolism. These enzyme production processes are dependent on an effective protein secretion machinery, whose regulation and expression are controlled by various transcription factors, as previously described. Factors such as the Hap2/3/5 complex, involved in the positive regulation of cellulases, and the *creA* factor, a key repressor in carbon catabolic repression (CCR), suggest regulatory mechanisms strongly related to carbon availability. Therefore, analyzing these secretion mechanisms is essential to understand how *L. gongylophorus* regulates enzyme expression and modifies its metabolism in relation to specific culture conditions.

#### 3.2.1. Differential gene expression in vesicular traficking and cellular recycling

Figure 2 shows the gene expression of genes related to vesicular trafficking, post-translational modifications and reutilization of cellular components under the conditions analyzed. The *erv29* gene (g5700) can be seen, highly overexpressed (3.48) in the exponential phase with cellulose, suggesting a strong activation of COPII-mediated vesicular transport. This overexpression may be in response to the high production of enzymes necessary to degrade a less accessible substrate such as cellulose [20]. Genes such as pmt2 (g5408) and tvp38 (g5360) over-expressed during the stationary phase of growth on cellulose with values of 2.10 and 2.00, respectively, may be indicative of the activation of secondary metabolism. On the other hand, genes related to vesicular transport, such as sec2 (g6350) and get3 (g6489), significantly reduced their expression during the stationary phase on glucose and exponential growth on cellulose.

**Figure 2:**
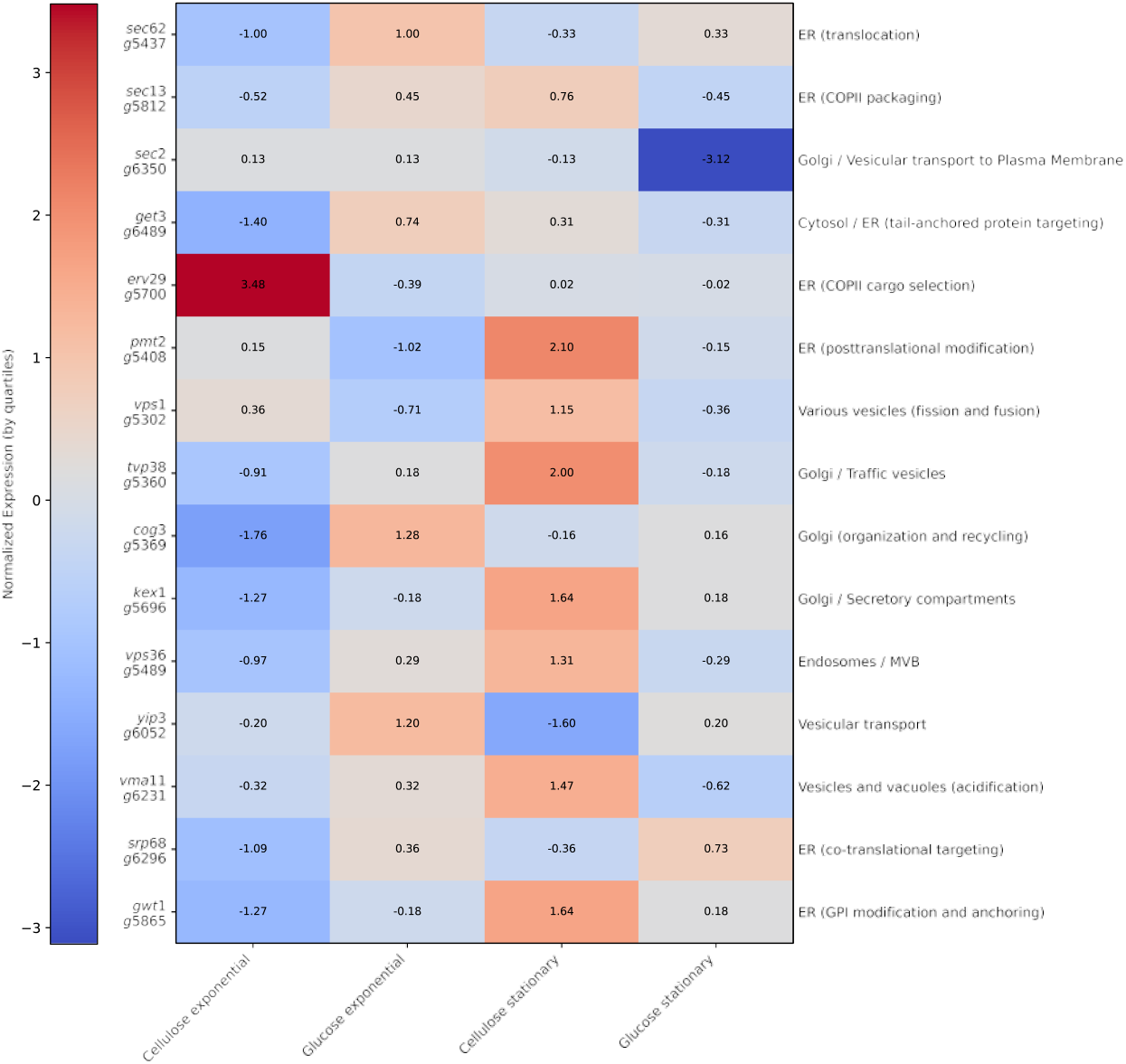
Heatmap illustrating differential gene expression (normalized by quartiles) of factors involved in vesicular trafficking, post-translational modifications, and cellular recycling under all experimental conditions. Color change represent relative expression levels: red indicates overexpression, blue indicates underexpression, and gray represents levels close to the mean. Annotations on the right side indicate the cellular localization and main functional role of each evaluated gene.

Starr et al. in 2018 investigated cellulase secretion in *Neurospora crassa*, demonstrating the dependence on specific endoplasmic reticulum (ER) cargo adapters such as *erv29* for efficient cellulase export [45]. These results may validate overexpression of the *erv29* gene during the exponential growth phase in cellulose, when cellulase secretion is maximal. The detection of extracellular nutrients can significantly influence the regulation of secretory machinery in lignocellulolytic fungi. Carbon limitation stress can activate mechanisms such as carbon catabolic repression (CCR), which can influence protein secretion [7]. Given the high expression of CAZymes, FOLymes, and sugar transporters expressed by *Leucoagaricus*, it is important to understand the metabolic response depending on the substrate or growth phase. The results obtained demonstrate the fine-tuning of secretion mechanisms in fungi, reflected in the differential expression of genes related to vesicular trafficking as an alternative pathway for the secretion of signal peptides of CAZymes enzymes reported in *Leucoagaricus* [9]. These results highlight the plasticity of the secretory system and the regulation of specific factors such as *erv29, sec2*, and *pmt2*, which are crucial to optimize protein secretion without losing cell viability.

### 3.3. Transcriptional landscape

Of the total of 6,748 genes reported for the *L. gongylophorus* genome [9], 69.7% (4,704 genes) presented at least one positive alignment and were considered in the differential expression analysis. From this subset, 2,031 genes (43.2%) showed significant changes in their expression (adjusted p-value < 0.1). Among the differentially expressed genes, 1,088 (23%) presented a positive log_2_FC, indicating an upregulation, while 943 (20%) showed a negative log_2_FC, suggesting a downregulation. With about 45% of differentially expressed genes we can appreciate a robust differential expression pattern and suggests that there may be several biological processes related to a substrate change or a growth phase. Additional analyses, such as functional annotation review and GO term enrichment, may help to understand the most involved biological processes.

To refine the selection of differentially expressed genes, we applied more restrictive statistical parameters. Table 3 summarizes the number of up- and down-regulated genes across conditions, highlighting expression patterns under different carbon sources and growth phases.

**Table 3.** Total upregulated and downregulated genes found for each pairwise analysis performed via DESeq2. Parameters: padj ≤ 1e-16, log_2_FC > 1 for upregulated genes, log_2_FC < -1 for downregulated genes, and log_2_FC between 1 and -1 for Not Changed Genes.

| Pairwise Comparison | Upregulated Genes | Downregulated Genes | Not Changed Genes |
| --- | --- | --- | --- |
| Glucose vs Cellulose (Exponential Phase) | 83 | 107 | 4 |
| Glucose vs Cellulose (Stationary Phase) | 98 | 53 | 9 |
| Exponential vs Stationary Phase (Glucose) | 24 | 17 | 3 |
| Exponential vs Stationary Phase (Cellulose) | 37 | 22 | 6 |

A higher number of differentially expressed genes are observed when comparing the two substrates than when contrasting different growth phases (Table 3). The combination of padj ≤ 1 × 10^−16^ and log_2_FC > 1 is a powerful strategy for filtering differential expression results and ensuring that the genes identified as differentially expressed are both statistically significant and biologically relevant [5]. Based on these pairwise comparisons, we conducted a more detailed analysis of each condition in each growth phase, focusing specifically on the expression of CAZymes and FOLymes, centering our analysis on the results presented in Table 3.

#### 3.3.1. CAZyome and FOLyome expression

The highest number of differentially expressed genes (DEGs) was observed between glucose and cellulose during the exponential growth phase (Table 3). The annotation of genes under these specific conditions revealed a significant enrichment of genes encoding enzymes of the CAZymes group, which are associated with the degradation of complex polysaccharides, particularly cellulose (Table S2).

The highly represented group of glyoxal hydrolases (GHs) is responsible for the breakdown of complex polymers, fundamentally those belonging to the subgroup of cellulases (Table S2). In 2020, Hori et al. cultivated the insect-associated fungus *Daldinia decipiens oita* in different carbon sources and compared the expressed protein profiles, concluding that in all the substrates used, including glucose and cellulose, the most highly expressed group was the GH [19]. In general, cellulose degradation requires three types of enzymes: endoglucanases (EG: EC 3.2.1.4), cellobiohydrolases (CBH: EC 3.2.1.91) and *β*-glucosidases (BGL: EC3.2.1.21), which function synergistically [50]. The overexpression of these genes in the exponential phase is directly related to the consumption of cellulose as the only carbon source. The expression of genes such as g1755 (GH family 7) and g681 (GH family 6) in the exponential phase is directly related to the consumption of cellulose (Table S2). Additionally, the genes involved in the degradation of substrates such as xylan and pectin also exhibited differential expression in these substrates. Specifically, genes g2346 and g4614, which encode the enzyme endo *β*-1,4-xylanase, were downregulated in the glucose substrate during both growth phases, whereas they were upregulated in the carboxymethyl cellulose. Interestingly, the expression of the gene g2772, which encodes the enzyme exo-polygalacturonase, and it is involved in the extracellular degradation of pectin, was downregulated in the exponential growth phase with glucose but upregulated in the stationary phase. A gene set enrichment analysis (GSEA) of the main differentially expressed metabolic genes was also carried out, where enrichment with more than 40 carbohydrate metabolism sequences and, more specifically, enzymes with hydrolase activity of compounds can be observed with O-glycosidic bonds (Figure 3). Given the high expression of cellulase and glyoxal oxidase enzymes, the enrichment of CAZy-encoding genes occurred late during growth on cellulose (dos Santos Castro et al., 2014). The results of the functional enrichment performed are shown in Figure 3, demonstrating that there is enrichment of several biological processes and molecular functions. The most enriched classifications include carbohydrate metabolic process (gene ratio = 0.108, 29 test sequences, adjusted p-value = 2.86 × 10^−14^), transporters (Gene ratio = 0.101, 27 test sequences, Adjusted p-value = 4.08 × 10^−3^) and polysaccharide catabolism (Gene ratio = 0.067, 18 test sequences, Adjusted p-value = 3.52 × 10^−12^)

**Figure 3:**
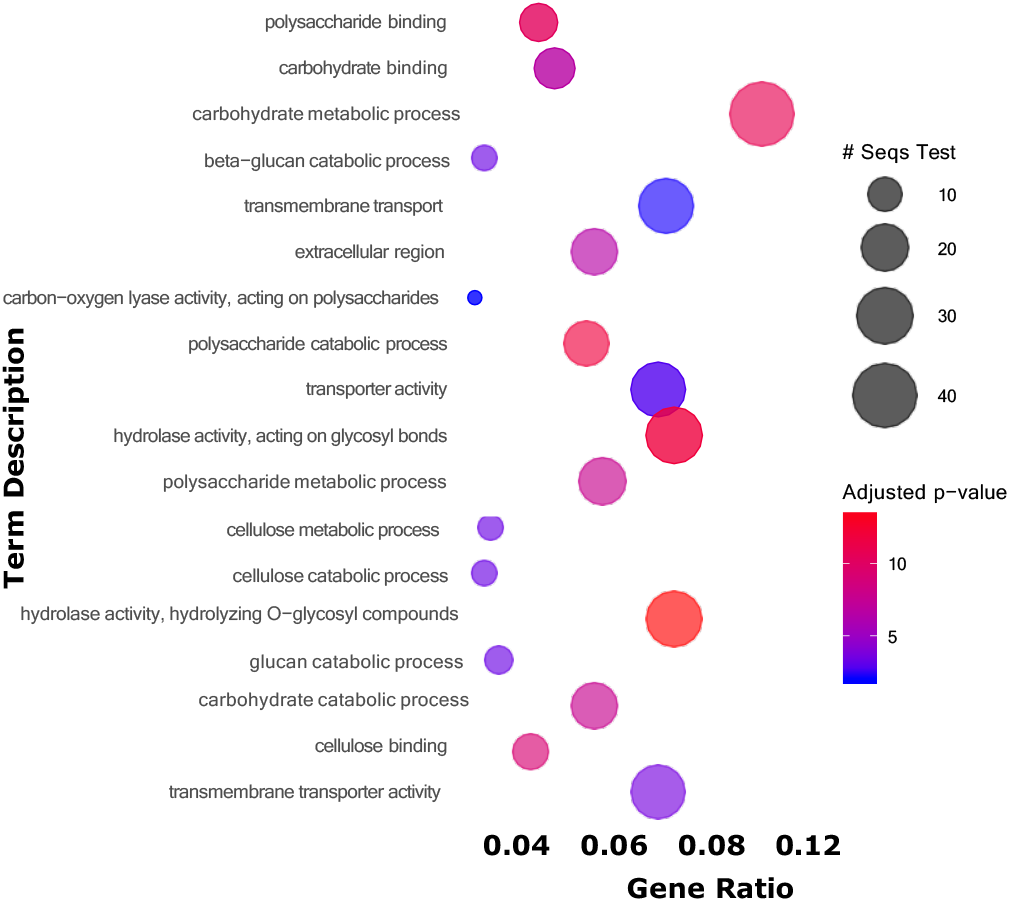
Bubble plot depicting the functional enrichment analysis performed. The y-axis shows the enriched functional annotation terms, while the x-axis represents the proportion of genes. The color gradient of the bubbles corresponds to the log10 of the adjusted p-value, with red indicating higher significance. The size of the bubbles represents the number of sequences associated with each term.

These results indicate that there is a high degree of specialization in the degradation and consumption of complex polysaccharides such as cellulose, as evidenced by the enrichment of hydrolase enzymes and transporters. In addition, enrichment of cellulose catabolism can be seen with a lower number of sequences but high significance (Gene ratio = 0.022, 6 test sequences, adjusted p-value = 1.19 × 10^−3^) Genes encoding upregulated CAZymes during growth on glucose versus cellulose are noteworthy, particularly those related to complex polysaccharide degradation that are typically suppressed under catabolic repression (CCR) [2][44].

### 3.4. Regulation of *creA* in relation to the expression of CAZymes depending on the substrate

Figure 4 shows a schematic representation of the expression mechanism of *creA* and CAZymes in fungi, specifically for *L. gongylophorus* according to the expression patterns obtained. According to the catabolic repression mechanisms reported, genes related to the degradation of complex polysaccharides such as cellulose should not be expressed (Figure 4, top left). Interestingly, during growth on glucose, genes such as g1024 (glycosyl hydrolase 18 family, pectinase), g4731 (glycosyl hydrolase 3, *β*-glucosidase), and g663 (glycosyl hydrolase 17, endo-*β*-1,3-glucanase) were overexpressed (Table S3), suggesting maybe constitutive expression [19].

**Figure 4:**
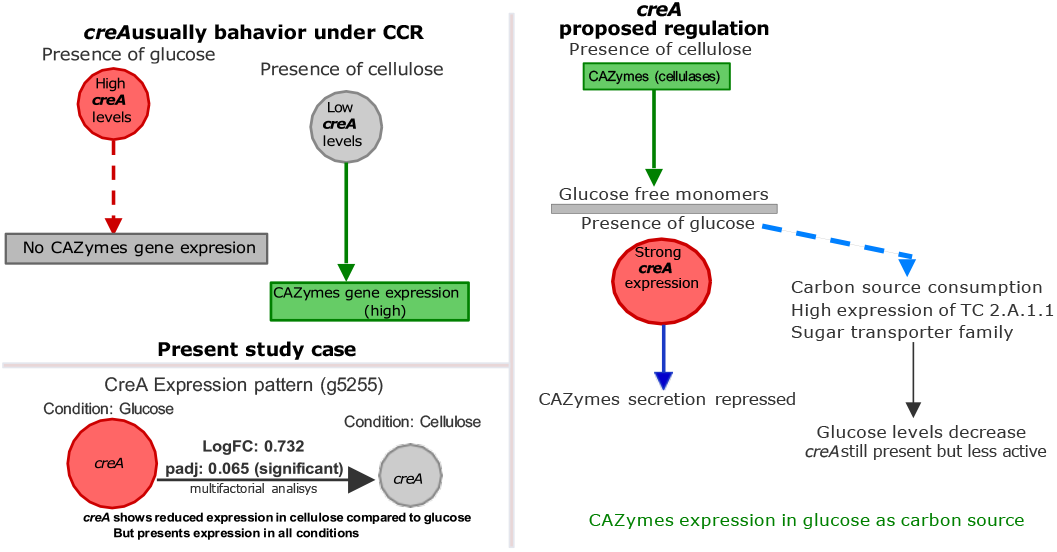
Schematic representation of the regulation of *creA* gene expression. On the top left, the most described mechanism of *creA* expression (CCR) is represented. On the bottom right, the expression pattern observed in this study is presented for all experimental conditions. Finally, on the left, a dynamic scheme of *creA* regulation in glucose and cellulose.

Previous studies have also reported instances of CAZymes expression not subject or capable to avoid catabolic repression. For example, dos Santos Castro et al. in 2014 observed 17 glycosyl hydrolases, including endoglucanase (*cel5b*) and *β*-1,4-glucanase (GH5), upregulated in *T. reesei* cultivated on glucose, cellulose, and sophorose [38]. Similarly, Henske et al. in 2018 documented high expression levels of GH enzymes during maltose (a glucose dimer) metabolism in *Pycnoporus cinnabarinus*, a white-rot fungus, while studying lignin degradation [18]. Conlon et al. in 2022 further supported these findings by demonstrating that *L. gongylophorus* expresses a full suite of enzymes required for cellulose degradation. Remarkably, their high-resolution in silico transcriptomic analysis revealed that these genes are expressed even in PDA medium without cellulose, indicating potential constitutive expression of biodegradative pathways [11]. According to the above, we propose a dynamic regulation scheme (Figure 4, right), where in the presence of cellulose, CAZymes (cellulases) are secreted and degrade the substrate. Due this hydrolysis process, degradation products are released and the expression of *creA* appears and regulate the secretion of CAZymes. This explains the levels of *creA* obtained in cellulose as a carbon source during exponential phase of growth. On the other hand, due to the consumption of glucose (or when it is the only carbon source), a large number of sugar transporters are activated that decrease the concentration of this monomer, the effect of *creA* decreased and enabling the expression of CAZymes in the presence of glucose. Peng et al. in 2021, presented a dynamic mechanism for regulating *creA* for Aspergillus niger when investigating the adaptation of this organism to low sugar concentrations. Initially, CAZymes degradation occurs at a low constitutive level, releasing sugars that are transported into the cell and activate specific TFs [34]. These findings collectively suggest that the expression of certain CAZymes genes in fungi, including *L. gongylophorus*, may be regulated either through constitutive mechanisms or influenced by catabolic repression, depending on the carbon source and growth conditions. Table 3 further shows the number of genes differentially expressed in glucose vs. cellulose during the stationary growth phase. In this comparison, the second largest number of DEGs was obtained. Under these conditions, genes related to cellulose degradation, such as cellulases A and C (g2688, g894 and g4409), are overexpressed (Table S4). In 2022, Datsomor et al. demonstrated cellulase activity in glucose and xylose [14], validating reports of the production of these enzymes by white rot fungi using glucose as the sole carbon source [8]. Other genes that are overexpressed in glucose during the stationary phase of growth are genes (g2573, siderophores; g920 and g1417, multicopper oxidase family; and g4536, *F e*^2+^ oxygenase). They are related to oxidative stress and the solubilization of Fe, and low availability of nutrients and Fe occurs in the stationary phase [52]. Recently, Schalamun et al. in 2023 demonstrated, while studying the regulatory protein RGS4 of *T. reesei*, that there is a relationship between the expression of siderophores and the expression of iron transporter genes during growth in glucose [39].

Many genes are downregulated in glucose during the stationary phase (overexpressed in cellulose in the stationary phase) (Table S4), with special attention given to those genes related to oxidative stress. Genes that are overexpressed in glucose to address oxidative stress have been previously described. The genes g5723 (BAG domain) and g2002 (small heat shock protein, HSP20) exhibit differential expression under the studied conditions (Table S4). Jain et al. (2018) reported that BAG genes are overexpressed in response to oxidative stress to prevent the induction of cell death and act synergistically with Hsp70 proteins. However, this synergy could not be demonstrated in *Aspergillus nidulans*. In our specific case, both genes are expressed under the same conditions, suggesting complementary roles in stress responses. Furthermore, Jain et al. proposed that BAG genes modulate secondary metabolism and sexual reproduction in fungi [21]. Based on our results, we infer that BAG expression may stimulate the production of secondary metabolites, such as trichodiene (g5783), in our biological model (Table S4). Additionally, two distinct strategies to cope with oxidative stress are observed, depending on the carbon source and growth phase. The modulation of secondary metabolism by BAG proteins likely occurs through multiple interconnected mechanisms. First, BAG proteins act as cochaperones by interacting with Hsp70 proteins to stabilize protein folding and prevent the accumulation of misfolded proteins [36][26]. This interaction not only protects the cell from oxidative stress but may also redirect cellular resources toward secondary metabolic processes, such as the biosynthesis of metabolites like trichodiene. Second, BAG proteins play a role in apoptosis regulation by inhibiting caspase activation and promoting cell survival under stress conditions. This function ensures cellular resilience, allowing fungi to maintain metabolic processes necessary for survival and development, including secondary metabolism and reproduction (Townsend et al., 2005). Lastly, BAG proteins may influence transcriptional regulation by interacting with transcription factors or signaling pathways that govern the expression of genes involved in secondary metabolism [21]. This could explain their role in modulating trichodiene production under specific conditions, as secondary metabolism is often tightly regulated by stress and environmental cues. In *L. gongylophorus*, the occurrence of such modulation would depend on the presence of oxidative stress-inducing conditions, functional Hsp70 proteins, and conserved signaling pathways linking BAG activity to secondary metabolism. While direct evidence for this mechanism in *L. gongylophorus* is lacking, studies in other fungal systems suggest that similar regulatory networks could be present, highlighting the need for further investigation. Given the low number of differentially expressed genes in the comparisons between exponential and stationary phase, both in glucose (24 up-regulated, 17 down-regulated, 3 unchanged) and cellulose (37 up-regulated, 22 down-regulated, 6 unchanged), we will continue the analysis with a global comparison of experimental conditions by analyzing the 50 genes with the highest differential expression.

As part of a multifactorial analysis across all pairwise analysis, Figure 5 presents a heatmap of the top 50 most differentially expressed genes among the analyzed samples. The upper part of Figure 5 shows the overexpression of laccases across all compared states. The expression of these FOLymes is significantly different when glucose is used as the carbon source, with the highest expression observed during the exponential growth phase. Notably, laccase was the only enzyme from the FOLymes group that showed differential expression in these substrates as the sole carbon source, highlighting a phase-dependent regulation of laccase production. Similar results were demonstrated by Hailei et al. (2015), who used the basidiomycete fungus Ganoderma lucidum to show that glucose is the most suitable carbon source for laccase production compared to other carbon sources such as glycerol, sucrose, lactose, starch, and corn flour [17]. Another possible explanation for these results is that the secretion pathways in the host organism are shared with endogenous cells [54]. Zhang et al. (2023) further demonstrated that the *cbh1* gene (exocellobiohydrolase) regulates the secretion of many enzymes, including laccases [56]. They also found that the expression of laccases increases when cellobiohydrolase is overexpressed during the exponential growth phase, particularly when glucose is used as the carbon source. This suggests that enhancing the activity of cellobiohydrolase in this phase can promote the production of laccases in *T. reesei*. Strong promoters, such as *pcbh1*, are highly expressed when cellulose is used as an inducer. While these promoters drive the production of various proteins, they also result in the production of endocellulases as byproducts, which can reduce the expression of target proteins, such as fusion proteins [35]. Additionally, in the presence of glucose as the sole carbon source, a broader range of polysaccharidases (pectinase, amylase, xylanase, and cellulase) is produced in *L. gongylophorus*, favoring the induction of a larger number of proteins [43].

**Figure 5:**
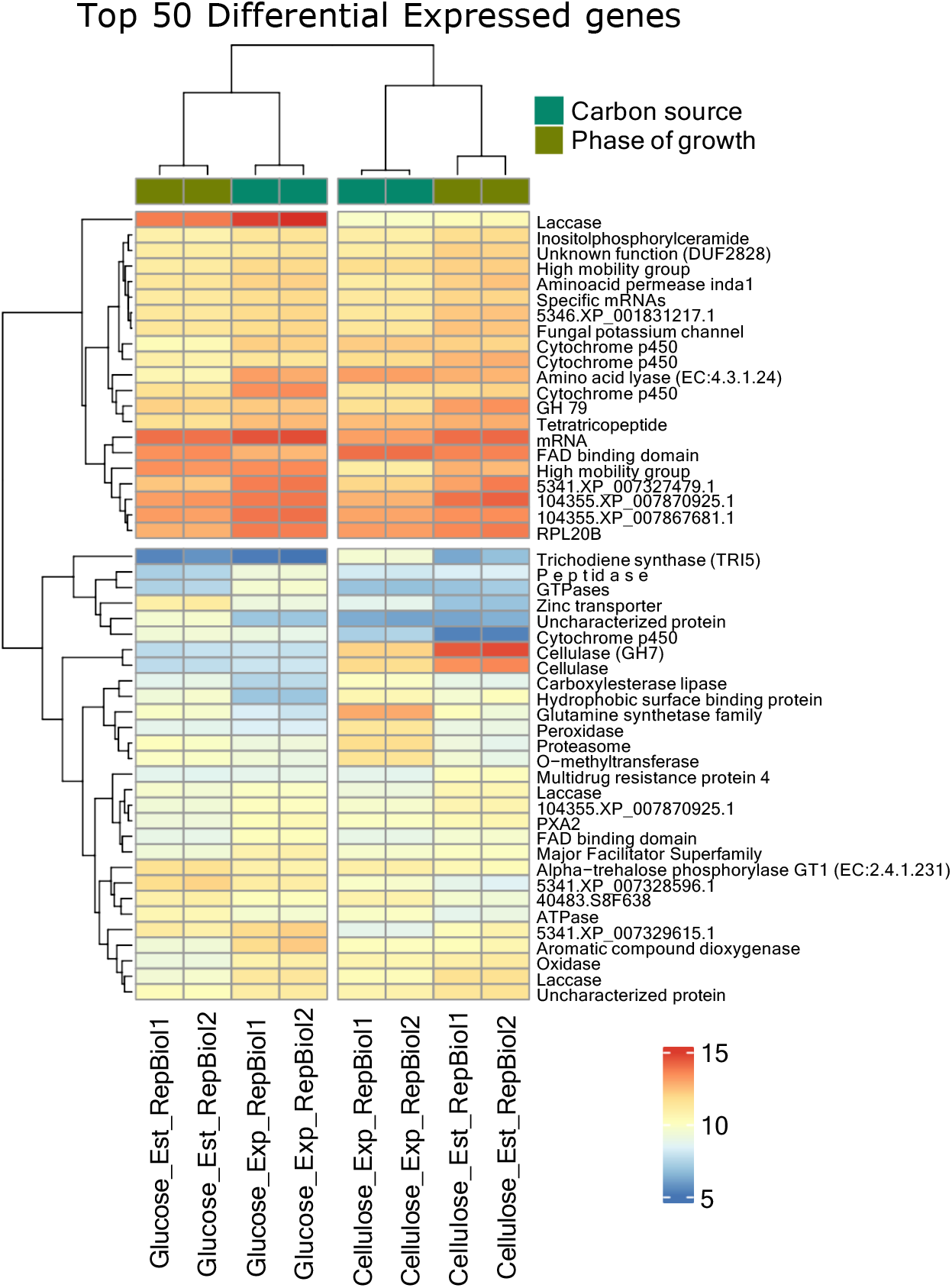
Changes in the expression of the 50 genes most differentially expressed genes among all the conditions analyzed. The change from blue to red indicates the change in the expression level.

There is also an overexpression of cellulases during the stationary phase of growth on cellulose, particularly of the GH7 group (cellulase family A). These enzymes are secreted into the medium to degrade cellulose. In Figure 2, overexpression of regulatory elements of the secretion apparatus was observed during this phase, thus correlating the phenomenon of these overexpressions in the stationary phase of growth on cellulose: a high production of cellulases leads to an increase in expression in the secretion pathways through the *pmt2* (g5408) and *tvp38* (g5360) genes.

The synthesis of trichodiene increases with cellulose degradation, and this correlates with the highest expression of trichodiene synthase during the exponential phase (Figure 5). This phase also coincides with peak levels of cytochrome P450 monooxygenase (CYP) expression, further emphasizing the important role of CYPs in polysaccharide metabolism and their potential involvement in regulating triterpenoid synthesis and other related metabolites. According to Zhou et al. (2021), the synthesis of triterpenoids is closely linked to carbohydrate metabolism, a key branch of cellular metabolism. These researchers demonstrated that increased polysaccharide catabolism provides more precursors for triterpenoid synthesis in the fungus *Ganoderma lucidum* [58]. This connection has been further supported by studies observing a decrease in polysaccharides such as chitin and *β*-glucan, which results in the inhibition of triterpenoid synthesis [55][51]. Additionally, reports have highlighted the regulation of terpenoid synthesis through modulation of the cytochrome P450 monooxygenase (CYP) family expression, suggesting that these enzymes play a crucial role in both terpenoid and carbohydrate metabolism. In the present study, four CYPs were differentially expressed among the top 50 most differentially expressed genes, with their expression consistently higher when cellulose was used as the carbon source. The differential expression of CAZymes and the increase in trichodiene synthesis during cellulose degradation, as observed in this study, highlights the intricate relationship between primary and secondary metabolism, particularly within the BCG clusters, which may regulate the synthesis of terpenoids like trichodiene.

### 3.5. Biosynthetic gene cluster expression

A total of 16 biosynthetic gene clusters (BGCs) were predicted from the L. gongylophorus genome encoding 8 terpene synthases (TSs), 4 NRPS-like fragments, 2 indole biosynthesis, 1 type I polyketide synthases (T1PKS) and 1 siderophore biosynthesis (Figure 6). For all the BGCs, positive alignments were found for the 4 conditions analyzed, although no differential expression was found for all the clusters, with only a few (Table S5).

**Figure 6:**
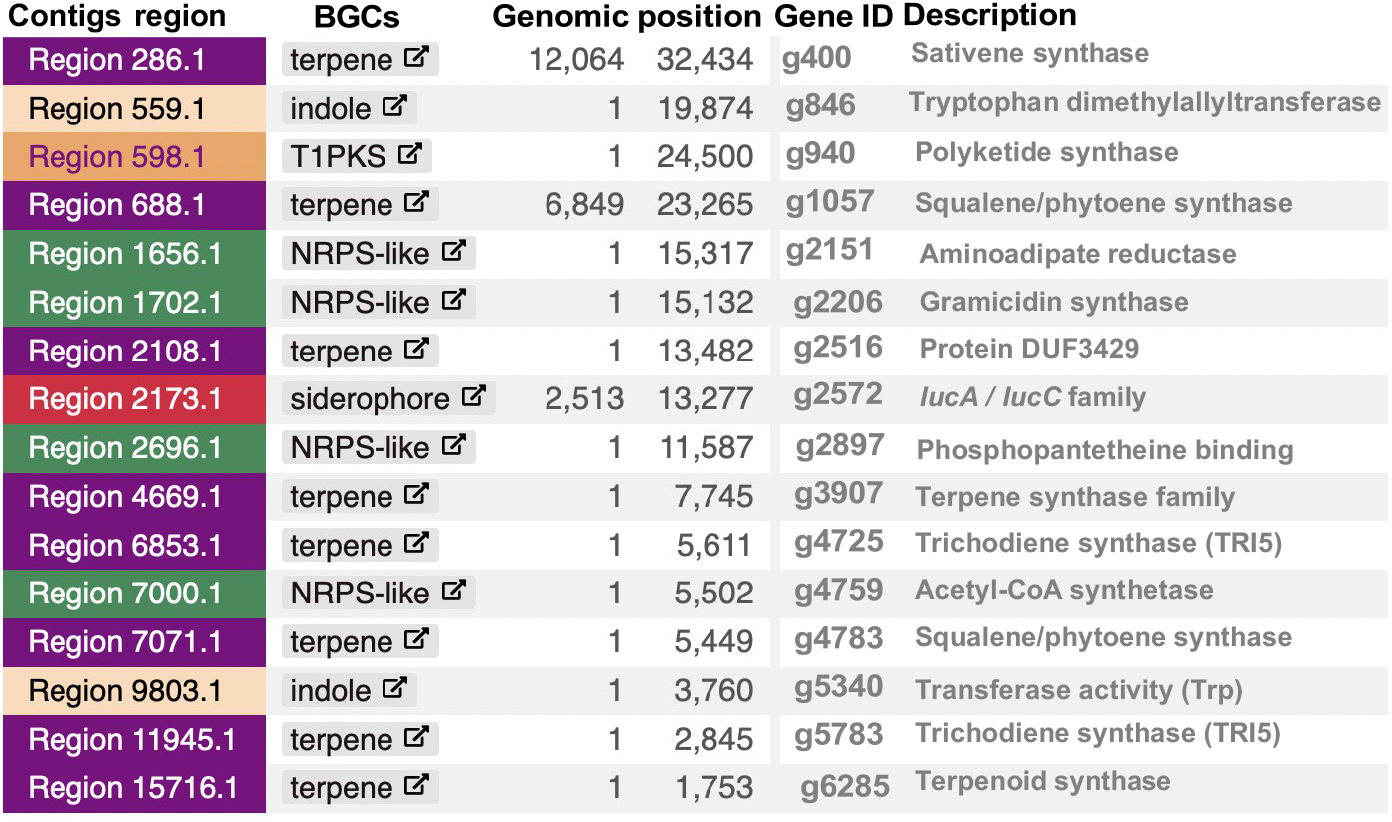
Biosynthesis-related gene clusters (BGCs) found in the *Leucoagaricus gongylophorus* LEU18496 genome and expressed in the transcriptomes. Only the BGCs are listed in the current figure. See Table S5 for the rest of the genes belonging to each cluster.

The production of terpenoids has been reported in several species belonging to the genera Ganoderma and Trametes, and their production has been evaluated in different fungi [1]. In 2022, Mao et al. studied terpene production in the white rot fungus *Ceriporia lacerata* during growth on cellulose [30]. In our case, we detected differential expression of trichodiene synthase (TRI5), which is encoded by the g5783 gene (Figure 6). This gene remains downregulated during the exponential phase and begins to increase in expression in the stationary phase of growth, which coincides with the increase in secondary metabolism when it does not compete with biomass precursors. However, the highest expression of TRI5 was observed during the exponential phase of growth in cellulose. Umezawa et al. reported that there is indirect evidence that positively regulated terpene synthase genes can favor the production of chelating agents in brown pudding fungi, especially if cytochrome P450 genes are expressed [48]. Although no DEGs related to the synthesis of siderophores were found, Figure 5 shows that DEGs were classified as cytochrome P450 genes. A large number of genes with cytochrome P450 functions were found to be positively regulated by both glucose and cellulose in both phases of growth. Although we are unable to confidently assign the exact functions of these genes, we can hypothesize, on the basis of the above results, that the overexpression of these genes favors secondary metabolism in *L. gongylophorus*. This relationship has been demonstrated in bacteria. Since in 2012, Lim et al. demonstrated the relationship between cytochrome P450 and siderophore production in bacteria [27].

## 4. Conclusions

As summary, the transcriptomic analysis showed that *L. gongylophorus* modulates its gene expression depending on the growth substrate and the growth phase. Principal component analysis confirmed that the type of substrate accounts for up to 99.5% of the variation in gene expression, indicating a clear segregation between experimental conditions. During the exponential phase, CAZymes and FOLymes are overexpressed when cellulose is used as the carbon source, facilitating the degradation of complex polysaccharides, whereas in glucose, a constitutive expression of CAZymes is observed, with variable expression of the creA gene to circumvent catabolic repression mechanisms; additionally, laccases are overexpressed in this carbon source, mainly during the stationary phase. During the stationary phase, significant metabolic differences were observed: in glucose, genes related to oxidative stress and nutrient release were activated, while in cellulose, secondary metabolism pathways, such as trichodiene production, were activated. These results underscore the importance of considering both the substrate and the growth phase to optimize the production of enzymes and other metabolites of industrial interest.

## Supporting information

All Supplementary Material

## Acknowledgements

We thank the CECIHTY for the PhD scholarship granted to Freddy Castillo Alfonso (CVU 943715) and for the projects CB-2016-287615 and A1-S-30750, which are essential funds granted for the development of this research. We thank Dr. Adrian Reyes Prieto from the University of New Brunswick for his valuable support and assistance during a research stay in his laboratory, which was fundamental for the transcriptomic data analysis. We also thank Ilse Salinas Peralta and UUSMB-UNAM for their technical assistance in RNA extraction and sequencing.

