## Supplementary material for "Gene expression analysis of carbohydrate catabolism in *Leucoagaricus gongylophorus* LEU18496": All Supplementary Material

[a]Posgrado en Ciencias Naturales e Ingeniería, Universidad Autónoma Metropolitana, Unidad Cuajimalpa, Av. Vasco de Quiroga 4871, Col. Santa Fe Cuajimalpa, Alcaldía Cuajimalpa, Ciudad de México, 05348, México[a]

[b]Department of Biology, University of New Brunswick 10 Bailey Drive, Fredericton, New Brunswick E3B 5A3, Canada[b]

[c]Departamento de Procesos y Tecnología, Universidad Autónoma Metropolitana, Unidad Cuajimalpa, Av. Vasco de Quiroga 4871, Col. Santa Fe Cuajimalpa, Alcaldía Cuajimalpa, Ciudad de México, 05348, México[c]

This PDF file includes:
Tables S1 to S5
Figs S1

**Table S1: STAR metrics obtained during multiple alignment process versus reference genome *Leucoagaricus gongylophorus* LEU18496.**

| | Cel_Est_RB1 | Cel_Est_RB2 | Cel_Exp_RB1 | Cel_Exp_RB2 | Glu_Est_RB1 | Glu_Est_RB2 | Glu_Exp_RB2 | Glu_Exp_ReB2 |
|---|---|---|---|---|---|---|---|---|
| **METRICS** | | | | | | | | |
| Reads | 10374785 | 10693803 | 15554611 | 14932036 | 8918017 | 9929675 | 11440257 | 12769133 |

| | | | | | | | | |
|---|---|---|---|---|---|---|---|---|
| Length | 151 | 151 | 151 | 151 | 151 | 151 | 151 | 151 |
| **UNIQUE READS:** | | | | | | | | |
| Mapped | 9601129 | 9895749 | 14361128 | 13748706 | 8295126 | 9119864 | 10519017 | 11729680 |
| % | 92.54% | 92.54% | 92.33% | 92.08% | 93.02% | 91.84% | 91.95% | 91.86% |
| **NUMBER OF SPLICES** | | | | | | | | |
| Splices | 4206986 | 5058526 | 6066012 | 5895863 | 3016364 | 4115421 | 4425385 | 5037089 |
| GT/AG | 4081613 | 4919806 | 5887934 | 5722987 | 2918254 | 3992009 | 4289396 | 4887271 |
| GC/AG | 109084 | 120112 | 153372 | 149424 | 79606 | 109068 | 111591 | 124563 |
| AT/AC | 761 | 1195 | 1160 | 1162 | 525 | 672 | 1010 | 1234 |
| Noncanonical | 15528 | 17413 | 23546 | 22290 | 17979 | 13672 | 23388 | 24021 |
| Mismatch rate | 0.40% | 0.44% | 0.43% | 0.41% | 0.40% | 0.42% | 0.47% | 0.46% |
| **DELETION** | | | | | | | | |
| Rate | 0.01% | 0.02% | 0.02% | 0.02% | 0.01% | 0.01% | 0.02% | 0.02% |
| Length | 2.79 | 2.81 | 2.79 | 2.83 | 2.64 | 2.78 | 2.7 | 2.64 |
| **INSERTION** | | | | | | | | |
| Rate | 0.01% | 0.01% | 0.01% | 0.01% | 0.01% | 0.01% | 0.01% | 0.01% |
| Length | 2.52 | 2.6 | 2.47 | 2.49 | 2.49 | 2.49 | 2.38 | 2.36 |
| **MULTI-MAPPING READS:** | | | | | | | | |
| Multiple loci | 133369 | 182784 | 196860 | 188895 | 110446 | 145777 | 172829 | 204261 |
| % Multiple loci | 1.29% | 1.71% | 1.27% | 1.27% | 1.24% | 1.47% | 1.51% | 1.60% |
| Too many loci | 652 | 2161 | 935 | 1012 | 476 | 844 | 1118 | 1831 |
| Too many loci | 0.01% | 0.02% | 0.01% | 0.01% | 0.01% | 0.01% | 0.01% | 0.01% |
| **UNMAPPED READS:** | | | | | | | | |
| Too short | 639630 | 613099 | 995687 | 993418 | 511966 | 663188 | 747282 | 833349 |
| % | 6.17% | 5.73% | 6.40% | 6.65% | 5.74% | 6.68% | 6.53% | 6.53% |
| Other | 5 | 10 | 1 | 5 | 3 | 2 | 11 | 12 |

Cel: Cellulose; Glu: Glucose; Est: Stationary growth phase; Exp: Exponential growth phase; RepBiol1 : Biological replicate 1 and RepBiol2: Biological replicate 2

**Table S2: Explained Gene ID based on annotations. Exponential phase of growth and contrast Glucose and Cellulose as carbon sources (Exp_Glc_vs_Cel)**

| Gene ID | Log2FC | pvalue | DEGs | Classification |
|---|---|---|---|---|
| g1000 | -3.599 | 1.63E-93 | DOWN | Meiotic DNA recombinase assembly |
| g1041 | -1.302 | 1.22E-16 | DOWN | Isocitrate lyase family |
| g1128 | -3.582 | 8.74E-41 | DOWN | Hydrophobic surface binding protein |

| | | | | |
|---|---|---|---|---|
| g1151 | -3.576 | 1.85E-22 | DOWN | AMP-binding enzyme C-terminal domain |
| g1271 | -2.125 | 3.66E-44 | DOWN | Belongs to the glycosyl hydrolase 28 family |
| g1282 | -1.982 | 8.98E-23 | DOWN | Glycoside hydrolase family 61 protein |
| g1301 | -2.480 | 1.78E-38 | DOWN | Alpha-galactosidase |
| g1369 | -2.272 | 3.53E-30 | DOWN | Starch/carbohydrate-binding module (family 53) |
| g1418 | -3.386 | 4.76E-42 | DOWN | Not anotated |
| g1419 | -2.601 | 3.30E-35 | DOWN | Belongs to the multicopper oxidase family |
| g1478 | -1.994 | 2.87E-29 | DOWN | AMP-binding enzyme C-terminal domain |
| g1506 | -3.323 | 5.19E-117 | DOWN | General substrate transporter |
| g1565 | -2.366 | 9.17E-84 | DOWN | Belongs to the MIP aquaporin (TC 1.A.8) family |
| g1601 | -2.507 | 2.32E-40 | DOWN | RNA cytidine acetyltransferase |
| g1602 | -5.990 | 1.58E-47 | DOWN | Beta-L-arabinofuranosidase, GH127 |
| g1702 | -1.621 | 2.86E-46 | DOWN | Belongs to the glycosyl hydrolase 31 family |
| g1736 | -1.997 | 2.44E-17 | DOWN | Sugar (and other) transporter |
| g1737 | -2.301 | 7.90E-51 | DOWN | Not anotated |
| g1755 | -8.395 | 0 | DOWN | Glycoside hydrolase family 7 |
| g1756 | -1.402 | 9.52E-27 | DOWN | Not anotated |
| g1787 | -2.351 | 3.74E-106 | DOWN | Belongs to the glycosyl hydrolase 5 (cellulase A) |
| g1793 | -1.672 | 4.74E-20 | DOWN | Major Facilitator Superfamily |
| g193 | -1.190 | 6.41E-27 | DOWN | Dehydratase family |
| g1955 | -2.387 | 5.13E-53 | DOWN | Glycosyl Hydrolase Family 88 |
| g2118 | -1.910 | 3.33E-19 | DOWN | Belongs to the cyclin family |
| g2167 | -1.018 | 1.00E-18 | DOWN | Aryl-alcohol oxidase |
| g2234 | -3.009 | 6.88E-20 | DOWN | Not anotated |
| g2259 | -1.716 | 9.52E-36 | DOWN | Pyridoxal-dependent decarboxylase |
| g2346 | -3.543 | 3.32E-173 | DOWN | Beta-xylanase |
| g2433 | -1.281 | 7.86E-21 | DOWN | Not anotated |
| g2441 | -1.500 | 4.74E-33 | DOWN | Not anotated |
| g2528 | -2.314 | 2.46E-24 | DOWN | O-methyltransferase |
| g2556 | -2.937 | 3.15E-50 | DOWN | Maceration and soft rotting of plant tissue |
| g2628 | -2.068 | 1.35E-76 | DOWN | Protein of unknown function (DUF1479) |
| g268 | -2.668 | 3.43E-107 | DOWN | Polysaccharide lyase family 4 protein |
| g2688 | -4.167 | 3.78E-137 | DOWN | Glycosyl hydrolase 7 (cellulase C) family |
| g2724 | -2.339 | 1.13E-57 | DOWN | 1,4-alpha-D-glucan glucohydrolase |
| g2756 | -1.456 | 4.30E-21 | DOWN | Not annotated |
| g2834 | -2.132 | 1.53E-26 | DOWN | Polysaccharide lyase family 4 protein |
| g2839 | -1.717 | 6.40E-37 | DOWN | MFS general substrate transporter |
| g2860 | -2.929 | 5.46E-18 | DOWN | Beta-galactosidase, domain 3 |
| g2876 | -2.577 | 5.32E-31 | DOWN | GMC oxidoreductase family |
| g2900 | -3.102 | 3.08E-153 | DOWN | Peroxidase, family 2 |

| | | | | |
|---|---|---|---|---|
| g3005 | -1.966 | 1.39E-25 | DOWN | Glycosyl hydrolase family 79 |
| g3011 | -5.024 | 1.64E-176 | DOWN | Sugar transporter (TC 2.A.1.1) family |
| g3039 | -1.475 | 7.32E-19 | DOWN | Pleckstrin homology domain |
| g3135 | -7.000 | 0 | DOWN | Sugar transporter (TC 2.A.1.1) family |
| g3136 | -4.102 | 6.29E-122 | DOWN | Alpha-L-arabinofuranosidase C-terminus |
| g3151 | -2.657 | 1.74E-39 | DOWN | Phosphoenolpyruvate phosphomutase |
| g320 | -1.762 | 3.57E-66 | DOWN | Sugar transporter (TC 2.A.1.1) family |
| g3395 | -1.220 | 6.69E-22 | DOWN | AMP-binding enzyme C-terminal domain |
| g3532 | -4.048 | 3.91E-84 | DOWN | Sugar transporter (TC 2.A.1.1) family |
| g3555 | -2.212 | 1.90E-36 | DOWN | gag-polypeptide of LTR copia-type |
| g3628 | -2.244 | 5.90E-27 | DOWN | Formate/nitrite transporter |
| g3693 | -1.620 | 7.96E-19 | DOWN | *Cys/Met* metabolism PLP-dependent enzyme |
| g3695 | -2.746 | 6.76E-32 | DOWN | Ferritin-like domain |
| g3730 | -3.224 | 2.13E-78 | DOWN | Not anotated |
| g3762 | -3.403 | 5.00E-49 | DOWN | Glycosyl hydrolase family 115 |
| g3769 | -1.607 | 6.12E-17 | DOWN | Glycosyl hydrolase family 65, N-terminal domain |
| g3874 | -1.828 | 7.99E-33 | DOWN | Fruit-body specific protein a |
| g3908 | -1.896 | 6.83E-19 | DOWN | Fungal specific transcription factor domain |
| g3914 | -3.518 | 1.74E-35 | DOWN | Not anotated |
| g4015 | -3.529 | 3.75E-148 | DOWN | GPI anchor biosynthetic process |
| g414 | -3.963 | 4.80E-96 | DOWN | OPT oligopeptide transporter protein |
| g4220 | -1.759 | 1.14E-21 | DOWN | not anotated |
| g4373 | -2.115 | 6.49E-34 | DOWN | Glycosyl hydrolase 12 (cellulase H) family |
| g4381 | -1.004 | 3.14E-22 | DOWN | Beta-L-arabinofuranosidase, GH127 |
| g4409 | -3.306 | 5.33E-203 | DOWN | Glycosyl hydrolase 5 (cellulase A) family |
| g4520 | -2.601 | 2.04E-50 | DOWN | Glycoside hydrolase family 74 protein |
| g4545 | -4.137 | 2.40E-141 | DOWN | PA14 |
| g4594 | -1.095 | 2.54E-20 | DOWN | Protein of unknown function (DUF4243) |
| g4614 | -3.107 | 5.31E-41 | DOWN | Beta-xylanase |
| g4683 | -2.576 | 4.33E-40 | DOWN | Basic region leucine zipper |
| g469 | -4.471 | 1.02E-96 | DOWN | Major Facilitator Superfamily |
| g470 | -2.112 | 6.22E-30 | DOWN | Glycosyl Hydrolase Family 88 |
| g4744 | -1.885 | 9.34E-17 | DOWN | Domain of unknown function (DUF1929) |
| g4825 | -1.649 | 3.87E-21 | DOWN | Not anotated |
| g4885 | -1.636 | 2.18E-17 | DOWN | Protein tyrosine kinase |
| g5184 | -2.909 | 3.66E-28 | DOWN | Phosphate transporter |
| g5318 | -3.445 | 7.02E-179 | DOWN | Fibronectin type III-like domain |
| g5387 | -5.226 | 6.32E-39 | DOWN | Pectate lyase |
| g554 | -1.959 | 5.20E-43 | DOWN | Major Facilitator Superfamily |
| g5568 | -3.471 | 3.73E-168 | DOWN | Sugar transporter (TC 2.A.1.1) family |

| | | | | |
|---|---|---|---|---|
| g5580 | -2.210 | 1.25E-23 | DOWN | Glycosyl hydrolases family 43 |
| g5590 | -3.834 | 1.54E-141 | DOWN | Fungal-type cellulose-binding domain |
| g5636 | -1.432 | 1.73E-16 | DOWN | Not anotated |
| g5638 | -4.708 | 4.20E-173 | DOWN | Cellulose-binding GDSL lipase acylhydrolase |
| g5640 | -1.996 | 5.30E-33 | DOWN | Cytochrome p450 |
| g5644 | -1.661 | 2.14E-21 | DOWN | Sugar transporter (TC 2.A.1.1) family |
| g5683 | -4.762 | 8.37E-55 | DOWN | Belongs to the glutamine synthetase family |
| g572 | -2.526 | 2.77E-21 | DOWN | Belongs to the glycosyl hydrolase 3 family |
| g5783 | -4.455 | 6.03E-55 | DOWN | Trichodiene synthase (TRI5) |
| g5926 | -2.325 | 2.43E-70 | DOWN | Phosphate |
| g6131 | -2.405 | 1.40E-48 | DOWN | Type-B carboxylesterase lipase family |
| g6144 | -1.900 | 3.99E-54 | DOWN | Major Facilitator Superfamily |
| g6154 | -1.350 | 9.02E-27 | DOWN | Helix loop helix domain |
| g6253 | -1.804 | 8.88E-19 | DOWN | Major Facilitator Superfamily |
| g626 | -3.616 | 1.21E-116 | DOWN | Belongs to the glycosyl hydrolase 1 family |
| g6422 | -1.875 | 2.09E-21 | DOWN | Sugar transporter (TC 2.A.1.1) family |
| g6526 | -1.444 | 2.77E-21 | DOWN | Endonuclease-reverse transcriptase |
| g681 | -4.025 | 6.22E-284 | DOWN | Belongs to the glycosyl hydrolase family 6 |
| g687 | -2.074 | 9.07E-27 | DOWN | Aminotransferase class I and II |
| g726 | -1.200 | 6.20E-22 | DOWN | Glycoside hydrolase family 15 protein |
| g800 | -1.920 | 2.09E-22 | DOWN | Cupin domain protein |
| g848 | -1.363 | 9.26E-43 | DOWN | FAD binding domain |
| g867 | -2.970 | 2.60E-51 | DOWN | Carbohydrate-binding module family 1 protein |
| g880 | -2.524 | 4.26E-39 | DOWN | Taurine catabolism dioxygenase |
| g896 | -1.167 | 1.02E-23 | DOWN | RBM39 protein |
| g978 | -1.158 | 5.91E-29 | DOWN | GAL4-like Zn(II)2Cys6 |
| g1024 | 1.385 | 6.15E-16 | UP | Belongs to the glycosyl hydrolase 18 family |
| g1033 | 1.453 | 1.78E-22 | UP | UTP--glucose-1-phosphate uridylyltransferase |
| g1210 | 2.361 | 9.78E-31 | UP | Belongs to the cytochrome P450 family |
| g1224 | 1.098 | 3.92E-23 | UP | Not anotated |
| g1227 | 1.555 | 2.78E-31 | UP | Belongs to the aldehyde dehydrogenase family |
| g1243 | 1.763 | 2.72E-20 | UP | Not anotated |
| g1248 | 1.851 | 5.09E-46 | UP | Nntiporter (CaCA) (TC 2.A.19) family |
| g127 | 1.121 | 2.25E-16 | UP | Not anotated |
| g1383 | 1.403 | 6.83E-23 | UP | Zinc finger |
| g1508 | 1.587 | 1.22E-16 | UP | Kinase-like |
| g1513 | 2.232 | 7.46E-39 | UP | Expressed protein |
| g1709 | 4.749 | 4.09E-33 | UP | Not anotated |
| g1710 | 1.299 | 3.79E-29 | UP | Aminotransferase class-V |
| g1741 | 1.981 | 3.68E-19 | UP | Not anotated |

| | | | | |
|---|---|---|---|---|
| g1819 | 3.597 | 6.84E-167 | UP | Belongs to the peptidase A1 family |
| g1857 | 1.537 | 4.15E-36 | UP | Not anotated |
| g1881 | 1.842 | 9.53E-30 | UP | GMC oxidoreductase |
| g1902 | 5.251 | 1.01E-270 | UP | Belongs to the multicopper oxidase family |
| g1903 | 4.735 | 4.23E-142 | UP | Iron permease FTR1 family |
| g1980 | 2.543 | 6.86E-40 | UP | ABC-2 family transporter protein |
| g2046 | 1.459 | 1.55E-17 | UP | Cytochrome p450 |
| g2144 | 1.318 | 1.05E-22 | UP | FAD binding domain |
| g2157 | 2.133 | 2.45E-34 | UP | Not anotated |
| g2237 | 1.540 | 2.16E-29 | UP | Protein of unknown function (DUF1308) |
| g23 | 1.091 | 7.87E-19 | UP | Glycolipid 2-alpha-mannosyltransferase |
| g2405 | 1.008 | 1.36E-18 | UP | HhH-GPD superfamily |
| g2476 | 1.568 | 5.19E-20 | UP | Survival protein SurE |
| g2565 | 4.302 | 1.62E-54 | UP | Endonuclease/Exonuclease/phosphatase family |
| g2662 | 1.840 | 8.46E-18 | UP | Mannose-1-phosphate guanylyltransferase |
| g2822 | 1.896 | 2.03E-30 | UP | Protein of unknown function (DUF1687) |
| g289 | 1.696 | 7.62E-28 | UP | Plasma membrane fusion involved in cytogamy |
| g2932 | 1.203 | 1.41E-23 | UP | Not anotated |
| g2955 | 1.242 | 2.23E-28 | UP | Belongs to the peptidase A1 family |
| g3021 | 2.238 | 8.78E-37 | UP | FAD-linked oxidoreductase |
| g3113 | 1.107 | 6.59E-20 | UP | Major Facilitator Superfamily |
| g3213 | 2.751 | 1.07E-43 | UP | Putative GTPase activating protein for Arf |
| g3408 | 1.499 | 5.33E-16 | UP | Not anotated |
| g3459 | 2.269 | 2.62E-30 | UP | Dehydrogenases family |
| g3627 | 1.282 | 1.74E-16 | UP | Not anotated |
| g3636 | 1.274 | 1.10E-18 | UP | Not anotated |
| g3679 | 1.269 | 1.19E-21 | UP | Phosphoprotein phosphatase |
| g3757 | 1.787 | 3.96E-23 | UP | 4-aminobenzoate hydroxylase |
| g3839 | 3.359 | 6.09E-22 | UP | Belongs to the peptidase S10 family |
| g3885 | 1.490 | 2.50E-43 | UP | mRNA, complete cds |
| g3886 | 1.612 | 2.42E-26 | UP | Carbohydrate-binding module family 19 protein |
| g409 | 1.434 | 1.74E-23 | UP | Belongs to the aldehyde dehydrogenase family |
| g4260 | 1.112 | 5.38E-25 | UP | Domain of unknown function (DUF1929) |
| g4367 | 2.320 | 1.16E-70 | UP | NADPH oxidase isoform 2 |
| g4403 | 1.850 | 1.59E-21 | UP | Fasciclin I |
| g444 | 1.248 | 2.52E-28 | UP | Zinc finger, C2H2 type |
| g4485 | 2.312 | 9.34E-26 | UP | Oxalate decarboxylase |
| g4494 | 1.163 | 3.53E-23 | UP | Belongs to the aldehyde dehydrogenase family |
| g4627 | 1.165 | 7.45E-17 | UP | Pro-kumamolisin, activation domain |
| g4695 | 3.422 | 4.68E-32 | UP | Peptide methionine sulfoxide reductase |

| | | | | |
|---|---|---|---|---|
| g4731 | 1.574 | 5.13E-35 | UP | Glycoside hydrolase family 3 protein |
| g4732 | 4.458 | 3.47E-17 | UP | Cerato-platanin |
| g4760 | 1.284 | 7.53E-34 | UP | Oxidation of uric acid |
| g4841 | 1.792 | 3.49E-27 | UP | G protein alpha subunit |
| g5054 | 2.627 | 3.52E-22 | UP | Alginate lyase |
| g5129 | 2.050 | 2.76E-47 | UP | Ferritin-like domain |
| g5211 | 3.502 | 9.63E-88 | UP | FAD-binding domain |
| g5268 | 2.130 | 2.28E-30 | UP | GMC oxidoreductase family |
| g527 | 3.197 | 6.56E-140 | UP | NADPH oxidase |
| g5302 | 2.420 | 4.89E-78 | UP | High mobility group |
| g5352 | 1.944 | 5.62E-21 | UP | Aromatic compound dioxygenase |
| g550 | 2.309 | 2.25E-19 | UP | DUF89 domain-containing protein |
| g5569 | 3.359 | 9.89E-40 | UP | Not anotated |
| g560 | 3.167 | 3.13E-42 | UP | Not anotated |
| g5632 | 1.158 | 2.67E-28 | UP | Type-B carboxylesterase lipase family |
| g5639 | 1.526 | 3.25E-43 | UP | Chitin deacetylase |
| g5654 | 1.989 | 4.30E-16 | UP | Chitinase |
| g5758 | 1.499 | 3.24E-17 | UP | Acid protease |
| g5984 | 2.776 | 1.48E-54 | UP | Ras family |
| g6037 | 1.544 | 2.12E-25 | UP | Not anotated |
| g608 | 1.555 | 7.43E-21 | UP | Heparinase II III family protein |
| g6101 | 1.983 | 3.90E-44 | UP | Cytochrome p450 |
| g6102 | 4.295 | 2.78E-123 | UP | Not anotated |
| g6146 | 2.233 | 3.02E-26 | UP | Broad-Complex, Tramtrack and Bric a brac |
| g6190 | 1.672 | 1.52E-21 | UP | Belongs to the cytochrome P450 family |
| g623 | 1.597 | 2.02E-31 | UP | Nucleoside triphosphate hydrolase |
| g6304 | 1.809 | 2.23E-29 | UP | Lysin motif |
| g6367 | 1.922 | 7.49E-53 | UP | Glycosyl hydrolase family 61 |
| g6462 | 1.214 | 9.25E-16 | UP | Not anotated |
| g6582 | 2.782 | 2.06E-37 | UP | Not anotated |
| g663 | 3.381 | 4.36E-27 | UP | Belongs to the glycosyl hydrolase 17 family |
| g756 | 1.763 | 1.72E-42 | UP | NADPH oxidase regulator |
| g790 | 1.222 | 1.32E-19 | UP | BUD22 |
| g920 | 1.068 | 3.87E-21 | UP | Belongs to the multicopper oxidase family |
| g931 | 1.263 | 1.07E-26 | UP | FAD binding domain |
| g947 | 3.173 | 3.27E-60 | UP | Tripeptidyl peptidase |

*Not Annotated: Genes: They were not noted during the functional annotation performed by Castillo-Alfonso et al in 2024

**Table S3: CAZymes expressed in glucose as a carbon source. Genes with positive alignments greater than 1000 and found for the 3 CAZymes databases and with confirmed EC number.**

| | | | | CAZy Families | |
|---|---|---|---|---|---|
| **Gene** | **EC#** | **EC name** | **HMMER** | **dbCAN_sub** | **DIAMOND** |
| g4718.t1 | 1.1.3.13 | alcohol oxidase | AA3_3(176-547) | AA3_e3 | *AA3_3 |
| g560.t1 | 1.14.99.- | GH synthase | AA14(1-256) | AA14_e1 | AA14 |
| g2236.t1 | 1.2.3.15 | Glyoxal oxidase | AA5_1(22-622) | AA5_e17 | AA5_1 |
| g5515.t1 | 1.2.3.15 | Glyoxal oxidase | AA5_1(20-550) | AA5_e2 | AA5_1 |
| g3016.t1 | 2.4.1.- | alpha-trehalose-phosphate synthase | GT24(1282-1441) | GT24_e1 | GT24 |
| g2780.t1 | 2.4.1.1 | glycogen phosphorylase | GT35(145-862) | GT35_e0 | **GT35 |
| g1215.t1 | 2.4.1.109 | Mannosyltransferase | GT39(71-312) | GT39_e12 | GT39 |
| g4009.t1 | 2.4.1.109 | Mannosyltransferase | GT39(63-302) | GT39_e9 | GT39 |
| g5408.t1 | 2.4.1.109 | Mannosyltransferase | GT39(79-317) | GT39_e30 | GT39 |
| g3500.t1 | 2.4.1.15 | alpha-trehalose-phosphate synthase | GT20(158-669) | GT20_e24 | GT20 |
| g1150.t1 | 2.4.1.16 | Chitin synthase | GT2(1235-1740) | GT2 | GT2 |
| g1574.t1 | 2.4.1.16 | Chitin synthase | GT2(167-342) | GT2 | GT2 |
| g4182.t1 | 2.4.1.16 | Chitin synthase | GT2(642-1161) | GT2 | GT2 |
| g724.t1 | 2.4.1.16 | Chitin synthase | GT2(191-353) | GT2 | GT2 |
| g1115.t1 | 2.4.1.18 | 1,4-alpha-glucan branching enzyme | GH13_8(231-527) | GH13_e20 | **GH13_8 |
| g6.t1 | 2.4.1.183 | 1,4-alpha-glucan branching enzyme | GH13_22(96-497) | GH13_e196 | GH13_22 |
| g1295.t1 | 2.4.1.231 | alpha-trehalose phosphorylase | GT4(460-618) | GT4_e3141 | GT4 |
| g254.t1 | 2.4.1.25\| | 4-alpha-glucanotransferase | GH13_25(1395-1753) | GH133_e0 | GH13 |
| g3072.t1 | 2.4.1.34 | 1,3-beta-glucan synthase | GT48(634-1403) | GT48_e0 | GT48 |
| g3468.t1 | 2.4.1.34 | 1,3-beta-glucan synthase | GT48(768-1540) | GT48_e0 | GT48 |
| g3412.t1 | 2.4.99.18 | glycosyltransferase | GT66(24-697) | GT66_e1 | GT66 |
| g219.t1 | 3.2.1 | alpha-amylase | GH13_5(161-345) | GH13_e5 | GH13_5 |
| g1024.t1 | 3.2.1.14 | endo-chitodextrinase | GH18(21-223) | GH18_e134+CBM5_e97 | GH18 |
| g1444.t1 | 3.2.1.14 | endo-chitodextrinase | GH18(7-352) | GH18_e20 | GH18 |
| g3885.t1 | 3.2.1.14 | endo-chitodextrinase | GH18(64-431) | GH18_e236 | GH18 |
| g2294.t1 | 3.2.1.145 | 1,3-beta-galactosidase | GH43_24(15-193) | GH43_e60+CBM35_e81 | GH43_24 |
| g2690.t1 | 3.2.1.20 | alpha-glucosidase | GH31_1(355-812) | GH31_e39 | GH31 |
| g4449.t1 | 3.2.1.20 | alpha-glucosidase | GH31_1(302-850) | GH31_e7 | GH31 |
| g2864.t1 | 3.2.1.21 | alpha-glucosidase | GH3(54-144) | GH3_e223 | GH3 |
| g3981.t1 | 3.2.1.21 | alpha-glucosidase | GH5_12(21-603) | GH5_e145 | GH5_12 |
| g1107.t1 | 3.2.1.22 | alpha-galactosidase | GH27(128-376) | GH27_e38 | GH27 |
| g3207.t1 | 3.2.1.24 | alpha-mannosidase | GH38(159-381) | GH38_e20 | GH38 |
| g3525.t1 | 3.2.1.28 | alpha-trehalase | GH37(81-652) | GH37_e7 | GH37 |
| g2724.t1 | 3.2.1.3 | Glucan 1,4-alpha-glucosidase | GH15(49-450) | GH15_e31 | GH15 |

*AA: Auxiliar activity, **GT: Glycosyltransferase, ***GHGlycoside Hydrolase

**Table S4: Explained Gene ID based on annotations. Stationary phase of growth and contrast Glucose and Cellulose as carbon sources (Esta_Glc_vs_Cel)**

| Gene | Log2FC | padj | DEGs | CLASSIFICATION |
|---|---|---|---|---|
| **g4158** | 6.0789 | 2.83E-97 | UP | Not anotated |

| | | | | |
|---|---|---|---|---|
| **g4888** | 3.9613 | 1.38E-24 | UP | Not anotated |
| **g5569** | 3.302 | 2.46E-37 | UP | Lytic polysaccharide monooxygenase* |
| **g2688** | 2.5303 | 4.28E-59 | UP | Glycosyl hydrolase 7 (cellulase C) |
| **g1672** | 2.3066 | 5.77E-33 | UP | Not anotated |
| **g1475** | 2.2912 | 1.51E-40 | UP | Not anotated |
| **g1741** | 2.0643 | 3.82E-20 | UP | Not anotated |
| **g2683** | 2.0387 | 1.64E-25 | UP | Pectate lyase |
| **g5867** | 2.0091 | 1.88E-32 | UP | PWI domain |
| **g894** | 1.9875 | 2.00E-28 | UP | Glycosyl hydrolase 5 (cellulase A) family |
| **g5926** | 1.9155 | 1.78E-52 | UP | Phosphate |
| **g4536** | 1.7107 | 8.34E-21 | UP | 2OG-Fe(II) oxygenase superfamily |
| **g5590** | 1.6854 | 9.44E-33 | UP | Fungal-type cellulose-binding domain |
| **g5273** | 1.666 | 5.43E-17 | UP | AAA domain |
| **g2259** | 1.6492 | 4.23E-36 | UP | Pyridoxal-dependent decarboxylase |
| **g1282** | 1.6344 | 2.49E-18 | UP | Glycoside hydrolase family 61 protein |
| **g1686** | 1.6129 | 5.92E-34 | UP | Glycosyl hydrolase family 79 |
| **g6147** | 1.5814 | 1.24E-22 | UP | Belongs to the multicopper oxidase family |
| **g2573** | 1.5806 | 1.34E-27 | UP | *IucA* / *IucC* family |
| **g3021** | 1.5647 | 2.14E-17 | UP | Oxygen-dependent FAD-linked oxidoreductase |
| **g5302** | 1.5642 | 6.22E-32 | UP | High mobility group |
| **g6139** | 1.5593 | 1.85E-16 | UP | KR domain |
| **g719** | 1.4418 | 1.73E-20 | UP | Ribosomal protein P0 |
| **g4545** | 1.4337 | 3.13E-20 | UP | Glycosyl hydrolase |
| **g1841** | 1.4241 | 1.54E-29 | UP | multidrug resistance protein 4 |
| **g4520** | 1.4037 | 4.08E-17 | UP | Glycoside hydrolase family 74 protein |
| **g5318** | 1.3984 | 1.48E-39 | UP | Fibronectin type III-like domain |
| **g920** | 1.3859 | 2.85E-34 | UP | Multicopper oxidase family |
| **g3135** | 1.3377 | 2.14E-17 | UP | Sugar transporter (TC 2.A.1.1) family |
| **g1156** | 1.3016 | 3.48E-23 | UP | NAD dependent epimerase dehydratase |
| **g1417** | 1.3008 | 5.85E-22 | UP | Belongs to the multicopper oxidase family |
| **g623** | 1.2057 | 2.67E-17 | UP | Nucleoside triphosphate hydrolase |
| **g5171** | 1.1625 | 6.11E-17 | UP | Monooxygenase |
| **g4409** | 1.1527 | 1.55E-27 | UP | Glycosyl hydrolase 5 (cellulase A) family |
| **g2772** | 1.1301 | 8.07E-22 | UP | Exo polygalacturonase (pectin degradation) |
| **g2751** | 1.0886 | 1.04E-33 | UP | Phosphate |
| **g4015** | 1.0801 | 3.12E-16 | UP | GPI anchor biosynthetic process |
| **g2346** | 1.0504 | 2.33E-18 | DOWN | Beta-xylanase |
| **g4539** | 1.0025 | 3.33E-18 | DOWN | Cytochrome P450 family |
| **g1677** | -1.0635 | 2.67E-17 | DOWN | Heat shock chaperonin-binding motif. |
| **g2562** | -1.123 | 4.61E-19 | DOWN | C-3 sterol dehydrogenase |

| | | | | |
|---|---|---|---|---|
| **g6164** | -1.2504 | 3.30E-17 | DOWN | Helix loop helix domain |
| **g5646** | -1.3338 | 5.68E-18 | DOWN | Not anotated |
| **g2610** | -1.379 | 8.19E-34 | DOWN | SAM-binding methyltransferase superfamily |
| **g3387** | -1.4511 | 1.12E-34 | DOWN | Not anotated |
| **g5807** | -1.4923 | 3.24E-17 | DOWN | DnaJ C terminal domain |
| **g4677** | -1.6541 | 6.14E-26 | DOWN | Belongs to the cytochrome P450 family |
| **g2321** | -1.6907 | 1.64E-27 | DOWN | Not anotated |
| **g6083** | -1.7033 | 7.42E-22 | DOWN | HAD-like protein |
| **g6293** | -1.8526 | 3.36E-21 | DOWN | ZIP Zinc transporter |
| **g4442** | -1.8769 | 5.80E-18 | DOWN | Not anotated |
| **g1478** | -1.9776 | 3.22E-26 | DOWN | AMP-binding enzyme C-terminal domain |
| **g5580** | -2.0193 | 8.79E-19 | DOWN | Glycosyl hydrolases family 43 |
| **g2528** | -2.3407 | 5.38E-24 | DOWN | O-methyltransferase |
| **g5723** | -2.4087 | 3.79E-34 | DOWN | BAG domain |
| **g2900** | -2.4211 | 6.62E-97 | DOWN | Peroxidase, family 2 |
| **g4683** | -2.7838 | 1.17E-44 | DOWN | Basic region leucine zipper |
| **g2863** | -2.9729 | 1.40E-109 | DOWN | BAG domain |
| **g5783** | -3 | 3.32E-31 | DOWN | Trichodiene synthase (TRI5) |
| **g5683** | -3.0737 | 3.99E-23 | DOWN | Belongs to the glutamine synthetase family |
| **g2002** | -3.8682 | 4.62E-24 | DOWN | Small heat shock protein (HSP20) family |

*Not Annotated: Genes: They were not noted during the functional annotation performed by Castillo-Alfonso et al in 2024

**Table S5: Total reads aligned with the genes belonging to the different BCG clusters listed below.**

| **g847.t1** | **g846.t1** | **g845.t1** | **g844.t1** | **g402.t1** | **g401.t1** | **g400.t1** | **g399.t1** | **g398.t1** | **g397.t1** | **Gene** |
|---|---|---|---|---|---|---|---|---|---|---|
| 8087 | 1002 | 3116 | 571 | 4189 | 1559 | 321 | 789 | 3915 | 2715 | **Cel_Exp_1** |
| 7657 | 983 | 2971 | 536 | 4229 | 1528 | 323 | 795 | 3869 | 2838 | **Cel_Exp_2** |
| 2921 | 317 | 2453 | 592 | 2751 | 907 | 192 | 656 | 2327 | 2107 | **Glu_Exp_1** |
| 3341 | 340 | 2765 | 634 | 2697 | 911 | 175 | 814 | 2731 | 2495 | **Glu_Exp_2** |
| 3139 | 467 | 3514 | 369 | 2871 | 1051 | 137 | 577 | 1920 | 2086 | **Cel_Est_1** |
| 3353 | 293 | 2970 | 283 | 2468 | 655 | 71 | 719 | 2032 | 2885 | **Cel_Est_2** |
| 2144 | 336 | 1955 | 404 | 2051 | 845 | 198 | 501 | 1365 | 1022 | **Glu_Est_1** |
| 3151 | 471 | 2497 | 496 | 3053 | 1089 | 276 | 757 | 1837 | 1580 | **Glu_Est_2** |
| indole | | | | terpene | | | | | | **BCG** |
| Belongs to the cytochrome P450 family | Tryptophan dimethylallyltransferase | FAD binding domain | Thi4 family | Homocitrate synthase family | GHMP kinases C terminal | synthase | RNA polymerase Rpb5, C-terminal domain | WD40 repeat-like protein | Ras subfamily of RAS small GTPases | **Classification** |

| **g2150.t1** | **g2149.t1** | **g1058.t1** | **g1057.t1** | **g1056.t1** | **g1055.t1** | **g1054.t1** | **g1053.t1** | **g941.t1** | **g940.t1** | **g848.t1** |
|---|---|---|---|---|---|---|---|---|---|---|
| 1219 | 284 | 1240 | 281 | 270 | 722 | 717 | 3481 | 1562 | 4167 | 24788 |
| 1260 | 271 | 1176 | 310 | 260 | 734 | 678 | 3280 | 1435 | 3747 | 24048 |
| 769 | 170 | 893 | 131 | 148 | 463 | 330 | 2224 | 938 | 859 | 6482 |
| 828 | 174 | 1003 | 119 | 161 | 566 | 408 | 2256 | 1137 | 838 | 6526 |
| 756 | 147 | 912 | 164 | 90 | 645 | 440 | 2395 | 927 | 1484 | 12039 |
| 720 | 54 | 703 | 69 | 73 | 582 | 263 | 1266 | 534 | 553 | 8604 |
| 683 | 122 | 541 | 139 | 92 | 426 | 271 | 2624 | 475 | 1046 | 8056 |
| 858 | 164 | 654 | 157 | 122 | 528 | 371 | 3254 | 520 | 993 | 11208 |
| NRPS-like | | terpene | | | | | | T1PKS | | |
| Alcohol dehydrogenase GroES-like domain | NUC153 domain | Vacuolar ATPase | Squalene /phytoene synthase | Not assigned | Not assigned | Aldo/keto reductase family | Provides the precursors necessary for DNA synthesis. | GAL4-like Zn(II)2Cys6 | polyketide synthase | FAD binding domain |

| **g4724.t1** | **g3907.t1** | **g3906.t1** | **g2897.t1** | **g2573.t1** | **g2572.t1** | **g2516.t1** | **g2515.t1** | **g2206.t1** | **g2205.t1** | **g2151.t1** |
|---|---|---|---|---|---|---|---|---|---|---|
| 712 | 4715 | 951 | 576 | 910 | 724 | 1321 | 14060 | 2530 | 2881 | 1416 |
| 737 | 4701 | 934 | 542 | 886 | 720 | 1271 | 13638 | 2603 | 2679 | 1392 |
| 874 | 2426 | 602 | 257 | 736 | 512 | 831 | 12814 | 861 | 841 | 882 |
| 997 | 2539 | 582 | 258 | 921 | 567 | 824 | 16063 | 765 | 831 | 963 |
| 772 | 4073 | 726 | 468 | 1779 | 534 | 928 | 7092 | 1695 | 2890 | 1079 |
| 794 | 2680 | 363 | 190 | 1137 | 417 | 622 | 6034 | 779 | 1660 | 582 |
| 465 | 3655 | 855 | 285 | 1001 | 393 | 459 | 5153 | 1430 | 954 | 940 |
| 675 | 4519 | 1132 | 335 | 1204 | 580 | 625 | 7499 | 1670 | 1210 | 1192 |
| terpene | terpene | | NRPS-like | siderophore | | terpene | | NRPS-like | | |
| synthesis of acetylglutamate | Terpene synthase family, metal binding domain | Major Facilitator Superfamily | Amp-dependent synthetase and ligase | lucA / lucC family | lucA / lucC family | Protein of unknown function (DUF3429) | ATPase (P-type) (TC 3.A.3) family. Type IV subfamily | Phosphopantetheine attachment site | Na h exchanger | Condensation domain |

| **g6285.t1** | **g5783.t1** | **g5430.t1** | **g4784.t1** | **g4783.t1** | **g4759.t1** | **g4725.t1** |
|---|---|---|---|---|---|---|
| 1046 | 1105 | 194 | 455 | 2861 | 3444 | 3180 |
| 989 | 1008 | 182 | 435 | 2693 | 3320 | 3013 |
| 164 | 34 | 97 | 525 | 2998 | 2113 | 1269 |
| 243 | 32 | 98 | 661 | 3043 | 2372 | 1323 |
| 318 | 67 | 95 | 311 | 1264 | 1582 | 1504 |
| 174 | 72 | 53 | 888 | 1057 | 793 | 1381 |
| 53 | 30 | 59 | 266 | 1583 | 2282 | 738 |
| 75 | 49 | 82 | 350 | 2142 | 2937 | 867 |
| terpene | terpene | indole | terpene | | NRPS-like | |
| terpenoid synthase | Trichodiene synthase (TRI5) | Transferase activity (Trp) | Not assigned | Squalene/phytoene synthase | acetyl-CoA synthetase-like protein | Trichodiene synthase (TRI5) |

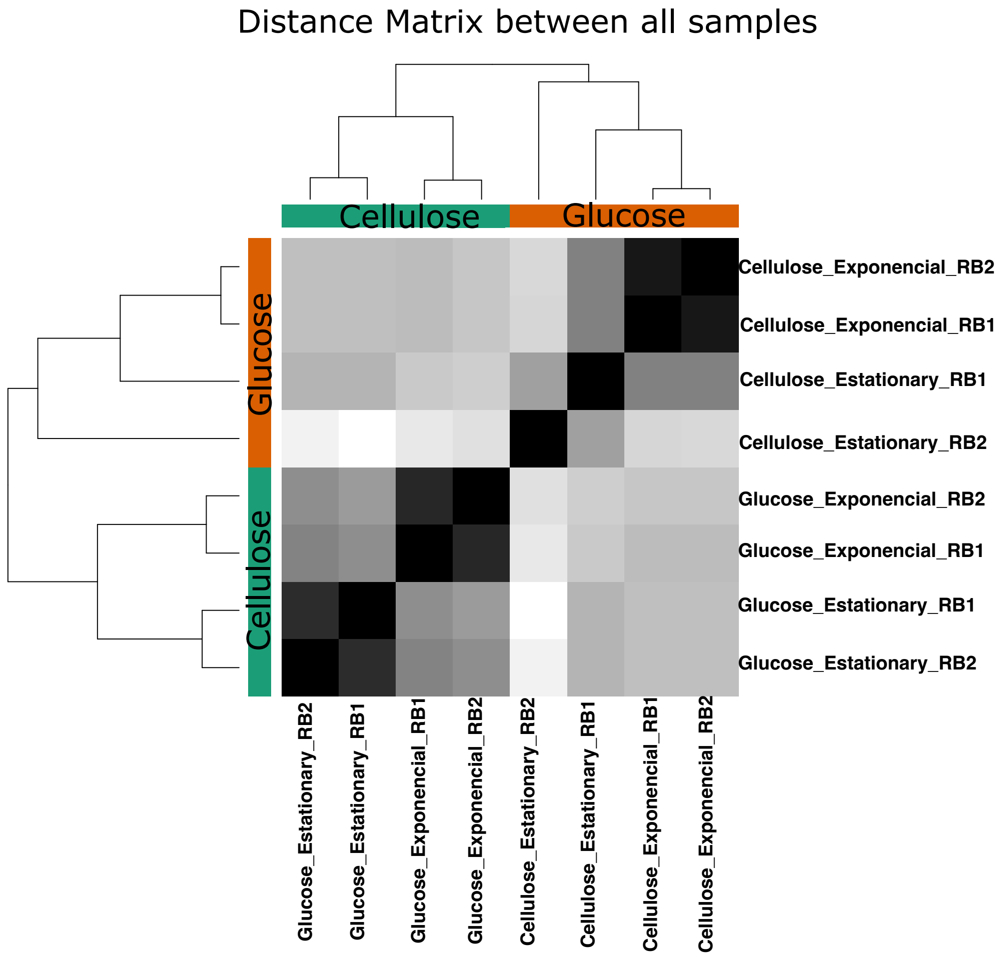


**Figure S1: Sample matrix distance of the 8 samples obtained using a distance matrix obtained using DESeq2 program in R language. RB1 and RB2 are the biological replicates. The intensity of the gray scale ranges from a value of zero representing the color white to 1 representing the color black.**
